# Architecture of a peptidoglycan peptidase complex involved in morphological transition in *Helicobacter pylori*

**DOI:** 10.1101/2025.08.29.672660

**Authors:** Olivier Danot, Ariel Mechaly, Aline Rifflet, Mathilde Bonis, Thimoro Cheng, Ahmed Haouz, Ivo Gomperts Boneca

## Abstract

Peptidoglycan is a meshwork macromolecule, made of polysaccharide strands cross-linked by short peptides, which encases the cytoplasmic membrane of bacteria and protects them against turgor pressure. Peptidoglycan peptidases are membrane or periplasmic enzymes that cleave these peptides, either lowering the cross-linking level of peptidoglycan to sculpt bacterial shape or allowing cell elongation by making space for the insertion of neosynthesized glycan strands. In the pathogen *Helicobacter pylori*, shape is important for virulence, and transition to a coccoid form after prolonged growth enables immune evasion. One particular endopeptidase, HdpA, is known to be involved in the maintenance of cell shape and in the transition to coccoids. Here we show that along growth, HdpA is constantly associated with LhiA, an inner membrane chaperone lipoprotein that keeps it in check while protecting it from fast proteolysis. The crystal structure of the HdpA-LhiA complex suggests that this interaction freezes the autoinhibitory interaction between the first domain of HdpA and the third, catalytic domain. Analysis of the evolution of the HdpA and LhiA protein levels over growth suggests that transition to coccoids is not triggered by a burst in HdpA activity but rather by a gradual weakening of the sacculus caused by the small fraction of free HdpA in equilibrium with LhiA-sequestered HdpA.

**SIGNIFICANCE STATEMENT:** The cell wall, the exoskeleton of bacteria, is the target of numerous antibiotics. Its principal component, peptidoglycan, is remodeled by an interplay between peptidoglycan synthases and hydrolases to accomodate bacterial growth. Because their activity can be harmful, hydrolases have to be tightly regulated. We discovered the dedicated inhibitory chaperone of a *H. pylori* peptidoglycan hydrolase and showed how the chaperone keeps the hydrolase in check in the cytoplasmic membrane and releases it slowly to allow it to perform its task, ultimately triggering a shape transition to a spherical form important for immune evasion. The crystal structure of the complex gives clues to the mechanism of peptidase activation, suggesting strategies to design agonists that could be used as antibacterials.

## INTRODUCTION

The bacterial cell wall is an essential, sturdy but elastic structure surrounding the cytoplasmic membrane of nearly all known bacteria. It acts as a mechanical and chemical barrier against external aggressions and internal turgor pressure. Its main component, peptidoglycan (PGN), is a meshwork-like macromolecule made of polysaccharide strands of alternate N-acetyl glucosamine (GlcNac) and N-acetyl muramic acid (MurNac) units cross-linked by short peptides(1, 2). PGN precursors are synthesized in the cytoplasm as MurNac saccharides with a pentapeptide stem whose typical sequence is L-Ala-D-Glu-2,6-diaminopimelic acid (mDAP)-D-Ala-D-Ala (with variations such as replacement of the mDAP by a lysine). These precursors are then conjugated to an undecaprenyl phosphate carrier and after addition of a GlcNac moiety, flipped into the periplasm, where they are incorporated into a nascent polysaccharide strand by glycosyl-transferase activities and cross-linked to the neighbouring strands by transpeptidase activities. The enzymes involved in glycosidic bond formation are either class A PBPs or SEDS proteins such as RodA and FtsW, while transpeptidation is ensured by class A and class B PBPs or L,D transpeptidases (3, 4).

In addition to these synthetases, bacteria also possess numerous enzymes involved in the cleavage of the different bonds present in the peptidoglycan: amidases hydrolyze the first amide bond between the MurNac and the peptide stem, peptidases hydrolyze the internal (endopeptidases) or C-terminal (carboxypeptidases) peptide bonds of the peptide stems or cross-links, and lytic transglycosylases cleave the glycosidic bond between MurNac and GlcNac in a reaction that generates a 1,6-anhydro bond in the MurNac. These enzymes fulfill extremely diverse functions, among which determining the shape and rigidity of the cell envelope as well as separation of daughter cells during cell division and making space for the insertion of new PGN strands in cell elongation (5, 6). Although the repartition of these tasks between the different enzyme classes is not strict, it seems that lytic transglycosylases and amidases, which break bonds irreversibly, have been co-opted for the definite process of daughter cell separation after division, while peptidases, whose activity leaves (at least for a subset of them) a residue able to play an acceptor role in the formation of a new cross-link, were selected for PGN remodelling processes involved in cell shape and elongation (6, 7). On the flip side, the activities of all of these enzymes, when not properly balanced by synthesizing activities, have the potential to degrade the PGN to a point of no return where the sacculus cannot contain the internal turgor pressure, resulting in spheroplast formation or lysis, hence their collective appellation of autolysins. Indeed, autolysins have been proposed to be involved in lysis upon treatment by antibiotics affecting PGN synthesis for more than fifty years (8), and their activities are controlled by numerous mechanisms, such as transcriptional regulation, proteolysis, protein-protein interaction (9), auto-inhibition (10, 11) or a combination of these. Although these regulations are increasingly studied, the molecular mechanism of the inhibition of endopeptidases only begins to be understood and structural information on the regulating complexes is still scarce.

The successful pathogen *Helicobacter pylori*, which colonizes half of the world human population, causing stomach ulcers sometimes leading to gastric cancer, has been shown to rely on its helical shape for its virulence (12). Furthermore, like other Campylobacterota (former Epsilonproteobacteria), it is able to undergo a morphological transition to a spherical (coccoid) form in response to different stresses or simply after prolonged growth (13). Although it is still unclear whether this morphology corresponds to a viable but non culturable form involved in persistence or transmission, the particular PGN composition of *H. pylori* or *Campylobacter jejuni* coccoids (14, 15) was found to make these pathogens invisible to the NOD1 innate immune receptor, likely contributing to the success of the infection, while a similar coccoid form elicited by the action of D,D endopeptidases was responsible for V. cholerae beta-lactam tolerance (16). In *H. pylori*, four peptidoglycan peptidases were found to be involved in the maintenance of cell shape: putative D,D endopeptidase Csd1, D,D endo/carboxypeptidase HdpA/Csd3, D,L carboxypeptidase Csd4 and L,D carboxypeptidase Csd6 (12, 17–19), with HdpA being the only one shown to be also involved in coccoid formation. Endopeptidase Csd1 is stabilized by its interaction with an enzymatically inactive homolog, Csd2, and an integral membrane protein, Csd7, forming a complex that is controlled by scaffolding protein CcmA (20). By contrast, nothing was known about the protein partners and localization of the HdpA endopeptidase. Here we examined how the activity, localization and stability of HdpA are regulated by a single lipoprotein and how this regulatory complex is involved in the transition to coccoids. Furthermore, we reveal the three dimensional architecture of the complex, suggesting a mechanism for the early stages of endopeptidase activation.

## RESULTS

### LhiA^S^ forms a stable complex with HdpA^S^ in vitro

HdpA is an enzyme involved in PGN remodeling and cell shape in *H. pylori*. In this bacterium, it is the only enzyme with a biochemically demonstrated peptidoglycan D,D-endopeptidase activity. Disruption of the *hdpA* gene causes cells to become slightly shorter and wider, as well as a change in PGN composition and the appearance of a subpopulation of multipolar, generally Y-shaped cells (12, 17). Furthermore, depending on the *H. pylori* strain, Δ*hdpA* mutations cause a delay (12) to a quasi-suppression (Fig. S1) of the transition towards the coccoid form, suggesting that HdpA plays a role in this transition.

The HP0762 protein of *H. pylori* 26695 strain (which we have renamed LhiA) is a predicted lipoprotein with a thioredoxin fold which was proposed to interact with HdpA on the basis of a yeast two-hybrid screen (21). It was therefore a candidate to be a new regulator of cell shape in this bacterium. We have set out to investigate this putative interaction and its effect on the enzymatic activities of HdpA. For that purpose, we improved HdpA purification to eliminate potential contaminating activities from *E. coli* peptidoglycan peptidase enzymes. This problem was previously bypassed by the use of penicillin G (17) but this strategy might represent a potential source of artefacts in the assays (e.g. titration of penicillin G by LhiA). Briefly, we used DV900(DE3), an *E. coli* strain lacking 8 low molecular weight PBPs (22) as a host to overproduce the HdpA protein devoid of its N-terminal hydrophobic segment (HdpA^S^ in the rest of the article), and we improved the purification protocol (Supplementary Material and Methods). The enzymatic activities of the obtained protein, measured in the absence of penicillin, were entirely inhibited by the addition of EDTA (Fig. S2A, B, C), indicating the absence of any detectable PBP contamination. Noteworthy, the effect of LhiA on these activities further suggests at most a very low level of contamination by *E. coli* peptidoglycan peptidases of any type (see below). LhiA devoid of its predicted signal peptide and lipidated cysteine (LhiA^S^) was purified in two steps also using the DV900(DE3) host and it was uncontaminated by peptidase activity (Fig S1B, F).

When separately subjected to exclusion chromatography on a Superdex 200 column, each of the LhiA^S^ and HdpA^S^ proteins eluted as a well-defined peak (Fig. 1A). The elution volume of LhiA^S^ corresponded to that of a globular protein of 20 kDa, as expected, whereas that of HdpA^S^ was that of a globular protein of 28 kDa, suggesting an interaction of HdpA^S^ (exact MW: 43.9 kDa) with the matrix of the column. When incubated together in stoichiometric amounts and subjected to exclusion chromatography on the same column, the peaks corresponding to the isolated proteins disappeared. They were replaced by a peak eluting earlier (apparent molecular weight 53 kDa), indicating the formation of a complex strong enough to survive the high salt (0.5 M NaCl) chromatography process and also interacting with the matrix. SDS-PAGE analysis of fractions collected around the elution peak confirmed the coelution of the two proteins.

**Figure 1:**
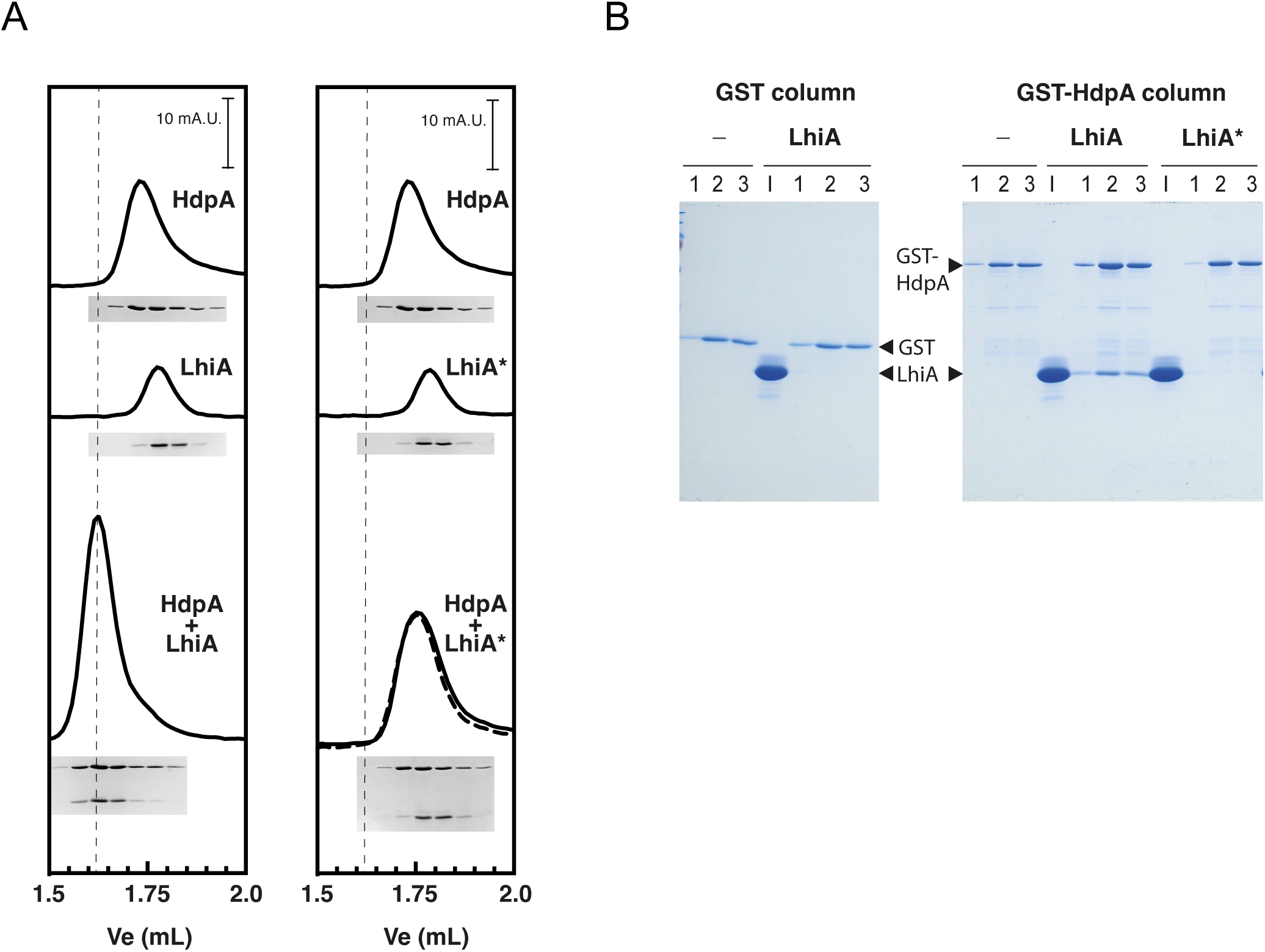
Interaction between LhiA and HdpA. A. Purified HdpA^S^ and LhiA^S^ or LhiA^S^* (10 µM) were preincubated for 15 min at room temperature either alone or together and 20 µL of the mix were injected on a Superdex 3.2 200 column equilibrated in buffer A. 280 nm UV traces are shown. The dotted vertical line marks the elution volume of the HdpA^S^-LhiA^S^ complex. The dotted trace on the right panel corresponds to the mathematical sum of the individual LhiA^S^* and HdpA^S^ traces. Since all filtration assays were performed in a row, the trace corresponding to HdpA^S^ alone is the same in the two panels. A Coomassie blue stained 13% polyacrylamide / bisacrylamide gel loaded with fractions positioned according to their elution volume is shown. In gels containing the two proteins, the HdpA^S^ band lies above the LhiA^S^ or LhiA^S^* band. B. Pull-down of LhiA by GST-HdpA. Purified LhiA^S^ or LhiA^S^* were run through glutathion-sepharose columns preadsorbed with GST or GST-HdpA. Equivalent volumes of LhiA^S^ input (I) and of three successive elution steps (1, 2, 3) were run on a 13% polyacrylamide / bisacrylamide gel which was stained with Coomassie Blue.

In a complementary experiment, purified GST (glutathion-S-transferase) or GST-HdpA^S^ fusion were immobilized on a glutathion sepharose column, and used for a pull-down assay with purified LhiA^S^ (Fig. 1B). LhiA^S^ was specifically retained on the GST-HdpA^S^ and not on the GST column, confirming the physical interaction between LhiA and HdpA.

### Structure of the HdpA-LhiA complex

The crystal structure of HdpA alone (23) showed that HdpA is a three-domain metalloprotein in which helix α3 of domain 1 occludes the catalytic cleft of the domain 3 by contributing through a glutamate 74 to the Zn^2+^ coordination sphere. Along with the inactivity of the crystallized protein, this suggested that HdpA was in an auto-inhibited conformation in the crystal. The auto-inhibitory nature of the interaction between domains 1 and 3 was clearly demonstrated in the case of the HdpA homolog ShyA (10).

To unravel the molecular details of the HdpA-LhiA interaction, we set out to crystallize the complex. Stoichiometric amounts of LhiA^S^ and HdpA^S^ were incubated together and the complex was further purified through exclusion chromatography. We obtained crystals that diffracted to 3.5 Å and the structure was solved by molecular replacement (Supplementary Table S1). In the crystal, the proteins appear to form a 1:1 complex, as suggested by the size exclusion chromatography experiments. Interestingly, the C-terminus of LhiA^S^ and the N-terminus of HdpA^S^ in the crystal were relatively close (Cα-Cα distance between K184^LhiA^ and Q36^HdpA^: 21 Å). To improve the resolution of the structure we decided to tether the two proteins in a way allowing the interaction observed in the complex. For that purpose, we constructed pOD-3, a pET24b(+) derivative encoding an LhiA^S^-SASG-HdpA^S^-LEHHHHHH hybrid protein. The protein was produced in BL21(DE3) and purified in three steps. We obtained crystals of the hybrid protein that diffracted to 2.09 Å (Supplementary Table S2).

As expected, the structure of the complex and that of the hybrid protein were extremely similar with an rmsd of 1.9 Å over 497 Cα, see Fig. S3). In the following, we will consider that the structure of the hybrid without the SASG linker represents that of the complex.

The structures of both proteins observed in the crystal (Fig. 2A) do not show any important conformational change with respect to the HdpA crystal structure or the LhiA Alphafold structure prediction (AlphaFold Protein Structure Database AF-O25457-F1) (23). The Cα rmsd for HdpA between the complex and the isolated protein is 1.79 Å over 356 Cα while the rmsd between the complex and isolated LhiA is 1.45 Å over 139 Cα. Notably, HdpA within the complex is in the auto-inhibited conformation described above. The interface between LhiA and HdpA is extensive (2000 Å^2^), consistent with the strong interaction observed in solution. LhiA mainly binds to auto-inhibitory domain 1 but also to domain 2 and the 1-2 linker, and it is located in the plane defined by the 3 domains of HdpA. The interface consists of a zipper of intricated secondary elements with, from bottom to top: β-sheet of HdpA domain 1, β7α4 C-terminal structure of LhiA, β7-β8 loop of HdpA domain 2, α1 helix of LhiA, β6 of HdpA domain 2 (Fig.2 B, C). Note that the interaction results in the extension of the β-sheet of HdpA domain 1 by the β7 strand of LhiA (Fig. 2C). The interface chemical nature is mixed, involving both polar/electrostatic and hydrophobic interactions with an interaction-dense region located at the turn of the β7α4 motif of LhiA. In this motif, E174 is in a position to establish hydrogen bonds to Q107 of HdpA domain 1, to the main chain of the linker between domain 1 and 2 and an electrostatic interaction with R186 of HdpA domain 2. Furthermore M175 is engaged in a knob-into-hole interaction (Fig. 2B) with HdpA Y86, L88, Q97 and L99 which protrude around it from the β-sheet of domain 1. It is clear that all these interactions between LhiA and HdpA domains 1, 2 and the linker inbetween must restrict the relative mobility between domain 1 and domain 2. It is also clear that these interactions take place in the context of the auto-inhibited form of HdpA. However whether LhiA interaction with HdpA stabilizes or destabilizes the auto-inhibited form remained to be determined.

**Figure 2:**
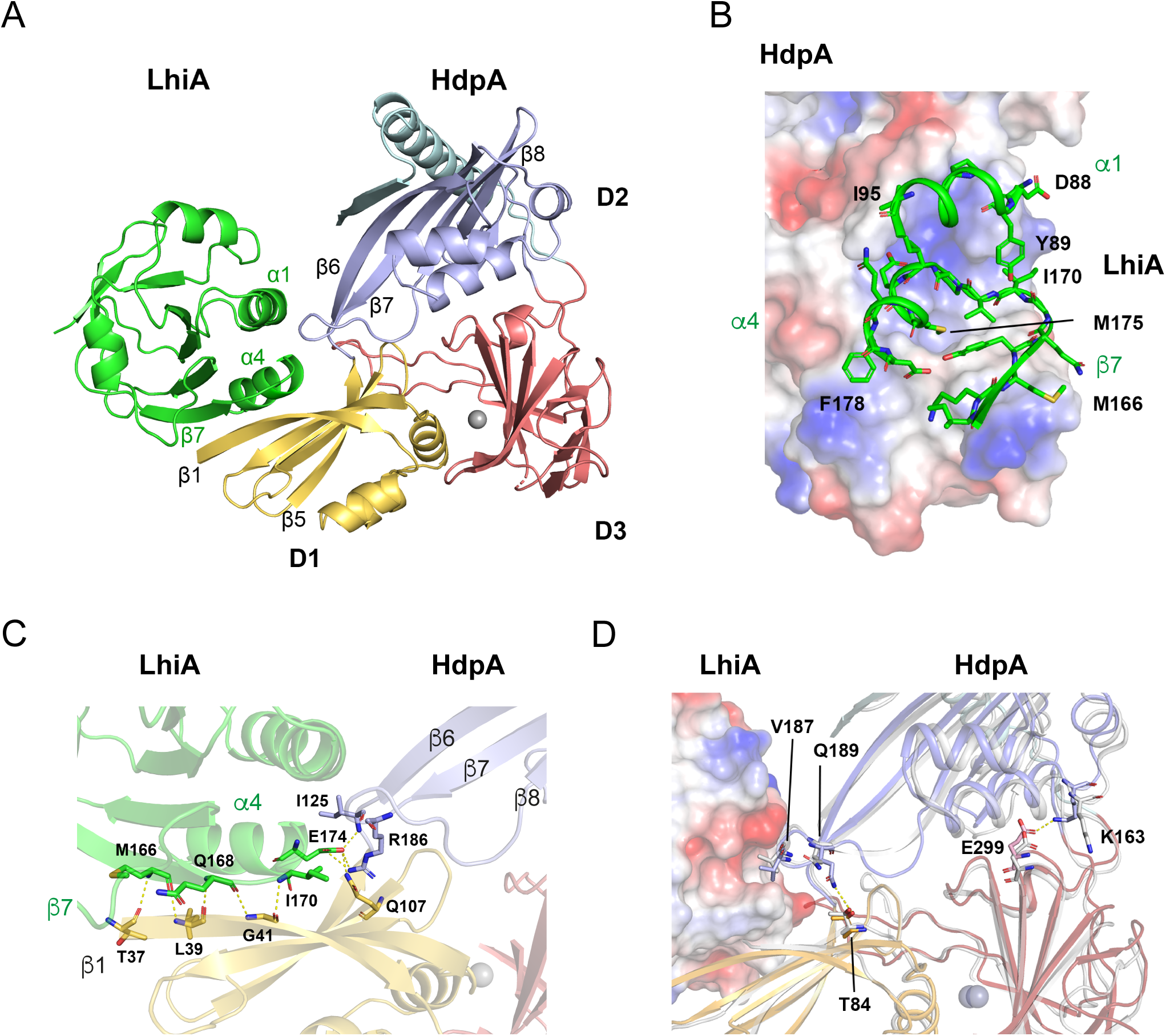
Three-dimensional structure of the HdpA-LhiA complex. A. Overall architecture of the complex with the three HdpA domains and the secondary structure elements (numbered as in An 2015 for HdpA) important for the interaction indicated. Color code will be conserved in the rest of the article.. B. HdpA surface potential with a cartoon representation of the secondary structure elements of LhiA involved in the interaction with HdpA showing residues (in particular M175) establishing polar and hydrophobic interactions. C. In the complex, the β7 strand of LhiA completes the β-sheet of HdpA domain 1 establishing typical hydrogen bonds between neighbouring mainchain carbonyl and amide groups. This positions E174 in a favorable position to establish hydrogen bonds and electrostatic interactions with HdpA Q107, R186 and I125. D. Two side-chains of HdpA residues establish domain domain interactions in HdpA only when it is complexed to LhiA. The LhiA-HdpA complex is shown (surface potential LhiA, cartoon representation of HdpA with color code of (A) superimposed with the structure of HdpA alone (PDB entry 4RNZ, in light grey). The movement of Q189 allowing it to interact with T84 could be elicited by the interaction between LhiA and residues R186 and V187 of the HdpA β7-β8 loop.

### The LhiA-HdpA interface in solution corresponds to that revealed in the crystal structure

Before answering that question, we wanted to validate in solution the LhiA-HdpA interface observed in the crystal. As a control we introduced in the expression plasmid used to produce LhiA two mutations predicted to affect this interface. These mutations result in the synthesis of a protein with a double E174R M175R amino acid substitution. The corresponding gene and protein will be noted *lhiA*\* and LhiA*, respectively, in the rest of the text. We purified the LhiA^S^* protein in the same conditions as LhiA^S^ and we tested the HdpA^S^-LhiA^S^* interaction by the same methods as for LhiA^S^. In the exclusion chromatography test (Fig. 1A, right panel), no interaction between LhiA^S^* and HdpA^S^ could be observed. Indeed when the two proteins were run together, the peak corresponding to the LhiA^S^-HdpA^S^ complex was absent, and replaced by a peak eluting earlier and corresponding to the sum (dotted trace) of the peaks obtained with the individual proteins. Note that due to the abnormal chromatographical behaviour of HdpA (see above), individual proteins eluted very close, which explains that only one peak is observed even though they are not interacting. Indeed analysis of the fractions by SDS-PAGE ((Fig. 1A, right panel, bottom gel) confirms that the two proteins do not coelute. Likewise, in the pull-down experiment, LhiA^S^* is not retained by the GST-HdpA^S^ column, contrary to LhiA^S^ (Fig. 1B). In conclusion, the E174R M175R substitutions targeting the interface observed in the crystal do abolish the HdpA^S^-LhiA^S^ interaction in solution, indicating that the crystal structure correctly recapitulates the architecture of the complex in solution.

### The HdpA peptidoglycan peptidase activities are strongly inhibited by LhiA

To understand the effect of LhiA on the conformation and activity of HdpA, we used the HdpA^S^ protein purified as described above in assays involving different types and concentrations of substrates. Soluble, HPLC-purified individual muropeptides were used as substrates to test different enzymatic activities of HdpA. The amount of substrate and product of a reaction were determined by HPLC analysis and quantification of the peaks corresponding to the different muropeptides (Fig.3 A, B, C and Fig S1A, C, D, E). To facilitate the analysis of the peaks, we used anhydromuropeptides, which had been shown to be substrates of HdpA in previous work (17). First, we reassessed the carboxypeptidase and endopeptidase activities of HdpA with purified HdpA^S^ (Fig. 3 D, E and Fig. S2) in initial velocity conditions (15% of the substrate consumed at most), using GlcNac-anhydroMurNac pentapeptide monomer (M5) and GlcNac-anhydroMurNac tetrapeptide dimer (D44), respectively. As expected from the fact that HdpA^S^ in solution exists mainly in its auto-inhibited form, the activities were low compared with that of *E. coli* MepA, a biochemically well studied LAS endopeptidase which does not seem to be auto-inhibited (24). Indeed a K_M_ could be assessed only for HdpA^S^ endopeptidase activity (120 µM, 95% confidence interval 90-160 µM, to compare with 13 µM for MepA). Carboxypeptidase activity of the pure protein was clearly much lower than endopeptidase activity, in line with the low amount of tetrapeptide in the *H. pylori* PGN composition. We also analyzed the composite activity of HdpA on GlcNac-anhydroMurNac tetrapeptide pentapeptide dimers (D54) at 8.5 µM. Interestingly HdpA hydrolyzed D54 in 3 different products: M5 and GlcNac-anhydroMurNac tetrapeptide (M4) which are the expected products of endopeptidase activity, but also D44 (Fig. 3B) which demonstrates that the carboxypeptidase activity of HdpA also acts on the acceptor stem peptide of muropeptide dimers, as had been proposed earlier (12). The composite rate of disappearance of D45, which is dominated by the endopeptidase activity, was similar to that of the endopeptidase activity measured on D44 (Fig. 3D), suggesting that as an endopeptidase HdpA does not discriminate importantly between pentapeptide-tetrapeptide and tetrapeptide-tetrapeptide cross-links.

**Figure 3:**
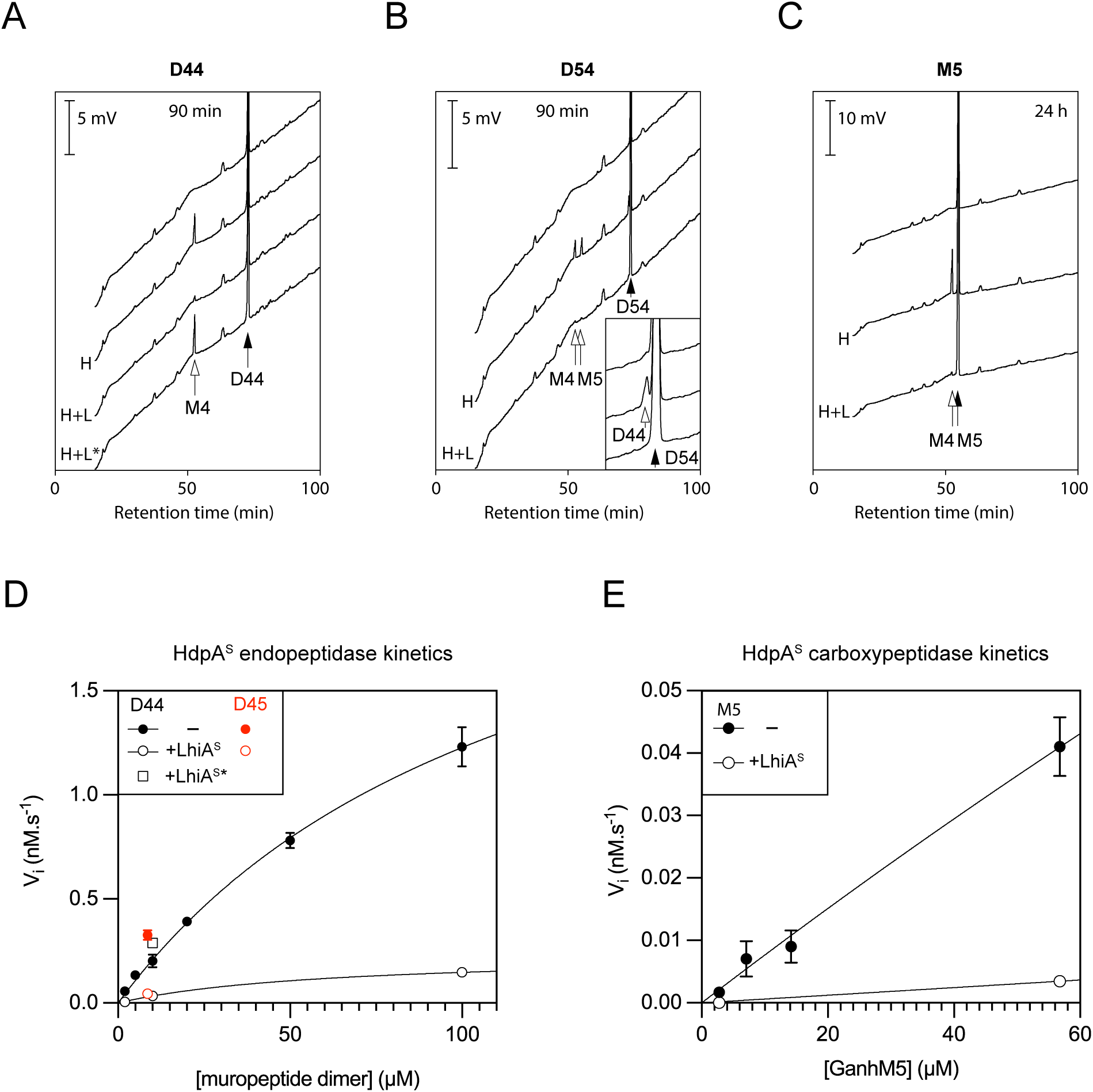
HdpA activities are severely inhibited by LhiA. Top panels: HdpA^S^ (0.4 µM) was incubated alone or in the presence of LhiA^S^ (0.4 µM) with the indicated muropeptide for the time indicated on top of the graph. Reaction products were separated by reverse phase HPLC on a Hypersil C18 Gold column as indicated in the Materials and Methods. Substrate (solid arrows) and products (open arrows) retention times are indicated.. H: HdpA^S^; H+L: HdpA^S^+LhiA^S^; H+L*: HdpA^S^+LhiA^S^*. A. Substrate : GanhM4-GanhM4 (D44), 10 µM. B. Substrate GanhM5-GanhM4 (D54) 8.5 µM. Inset: zoom on the 72 min retention time region showing the GanhM4-GanhM4 product. C. Substrate: GanhM5 (M5, 56 µM except top trace, 7 µM). D. Initial velocity of the endopeptidase reaction (disappearance of the substrate) as a function of substrate concentration for substrates GanhM5-GanhM4 and GanhM4-GanhM4, with and without LhiA^S^ or LhiA^S^*. E. Initial velocity of the carboxypeptidase reaction as a function of substrate concentration for substrate GanhM5. All experiments were performed at least in triplicate.

The effect of LhiA^S^ on all these activities was drastic: in a stoichiometric ratio with HdpA^S^, LhiA^S^ inhibited at least 10-fold the endopeptidase activity whether the substrate was D44 or D54 (Fig. 3A, B, D) at all anhydromuropeptide concentrations tested. It also inhibited the carboxypeptidase activity whether the substrate was M5 or D54 (Fig. 3B, C, E). The inhibition depended on an intact HdpA-LhiA interface since LhiA^S^* added at a 1.5:1 ratio did not decrease the HdpA^S^ endopeptidase activity on D44 (Fig. 3A, D). These results show that the interactions between LhiA^S^ and HdpA^S^ result in a complete shut-down of HdpA^S^ enzymatic activities. Along with the way LhiA is wedged between HdpA domain 1 and domain 2 in the crystal structure of the complex, they suggest that LhiA acts first by freezing domain 1 and domain 2 in the relative position they are occupying in the auto-inhibited state, which in turn defavors the opening of the structure.

This drastic and extremely specific effect of LhiA also confirms that the activities observed for the newly purified HdpA are genuinely due to HdpA and not to contaminants, which is an important issue while dealing with low activities frequently encountered with purified PGN remodeling enzymes.

### Deletion of the *lhiA* gene phenocopies the deletion of *hdpA*

Since LhiA^S^ inhibits the activities of HdpA^S^ in vitro, we expected that disrupting *lhiA* in vivo would result in an HdpA overproduction phenotype. To test this hypothesis, we constructed OMD1, a Δ*lhiA::aphA3* mutant derivative of the N6 strain as well as OMD2, an *lhiA\**-*aphA3* mutant which harbours the point mutations resulting in replacement of the LhiA interface residues E174 and M175 by arginine residues. We first followed their growth kinetics in liquid medium observed only a minor effect of Δ*lhiA* on growth (Fig. S4). We then examined their cell shape and their ability to form coccoids upon prolonged growth. Surprisingly, instead of mimicking an HdpA overproduction, both Δ*lhiA* and *lhiA*\* alleles recapitulate two prominent phenotypes of the Δ*hdpA* mutant, albeit in a less severe way. More precisely, Δ*lhiA* or *lhiA*\* were associated with the appearance of multipolar cells in exponential phase at a frequency ∼1/3 of that observed for Δ*hdpA* (Fig. 4A and S5A). Furthermore, these two mutations also resulted in a strong reduction in the proportion of coccoid forms generated after 120h growth, to a level close to that reached by the Δ*hdpA* mutant (Fig. 4B and S5B, C). The effects of Δ*lhiA* or *lhiA*\* mutations on cell width and length, however, were less clear than those observed with the Δ*hdpA* mutations (Fig. S6). Altogether, the overall positive genetic interaction between *lhiA* and *hdpA* suggested the possibility that LhiA protects HdpA from proteolysis.

**Figure 4:**
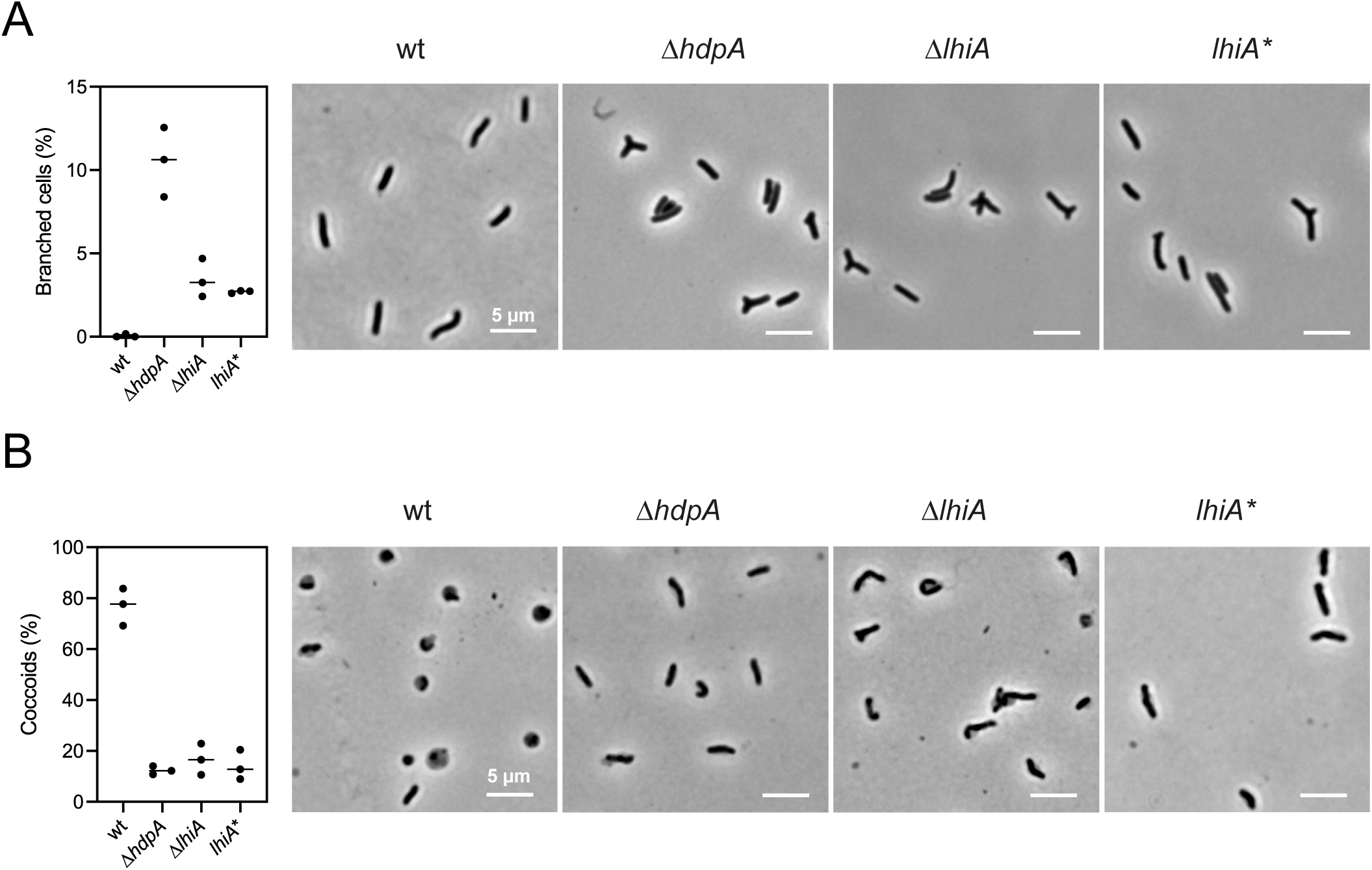
Δ*lhiA* and *lhiA\** mutations phenocopy Δ*hdpA* mutations for branched cell occurrence and coccoid formation, with a reduced severity. A. Cultures of strains N6, N6Δ*hdpA*::*aphA3*, OMD1 and OMD2 were grown in exponential phase (OD 0.3-0.8) and live cells were imaged by phase contrast microscopy. Left panel: Branched cells were identified by visual inspection of at least 450 cells from each of 3 independent cultures for each strain grown in exponential phase (OD 0.3-0.8). Individual results with confidence intervals are shown Fig. S5A. Right panel: representative phase contrast microscopy images of wt and mutant cells in exponential phase. B. Cultures of strains N6, N6Δ*hdpA*::*aphA3*, OMD1 and OMD2 were grown for 120h starting at OD 0.05 and live cells were imaged by phase contrast microscopy. Left panel: at least 300 bacterial cells from each of 3 independent cultures for each strain were identified using MicrobeJ and assigned to the rod or coccoid form as indicated in Material and Methods (individual results with confidence intervals in Fig. S5B, representative microscopy images in Fig. S5C). Right panel: examples of phase contrast microscopy images of rod cells (wt) and mutipolar cells obtained with the mutants.

### LhiA protects HdpA from proteolysis

To test this hypothesis, we examined the total amount of HdpA in the Δ*lhiA* and *lhiA*\* mutants (strains OMD1 and OMD2) by Western blot at the end of exponential phase and after prolonged growth because of the role of HdpA in coccoid formation (Fig. 5A). The most striking result of this experiment was that the HdpA protein was strongly depeleted in exponential phase in OMD1 and OMD2 compared to the wild-type strain. The fact that the *lhiA\** mutation gave a similar result as the *lhiA* deletion indicates that the LhiA-HdpA interface is involved in the effect of LhiA on HdpA. This suggests that this effect occurs at the post-translational level, in other words that LhiA protects HdpA from proteolysis by an unknown protease. In addition, the levels of HdpA decreased significantly after prolonged growth in all strains, so that HdpA levels become undetectable in the mutants, suggesting that the balance between synthesis and degradation of HdpA evolves in favor of degradation as growth proceeds. The fact that the Δ*lhiA* mutation abolishes both inhibitory and stabilizing activities of LhiA, along with the decreasing levels of HdpA along growth, provide an explanation for the complexity of the observed phenotypes: in exponential growth in the Δ*lhiA* context, HdpA levels are low but detectable and the activity of the enzyme is not inhibited, so that the total level of HdpA activity could be comparable to that in the wt. After prolonged growth however, HdpA becomes undetectable in the Δ*lhiA* context and late phenotypes such as the delayed transition to coccoids could develop more clearly than earlier phenotypes.

**Figure 5:**
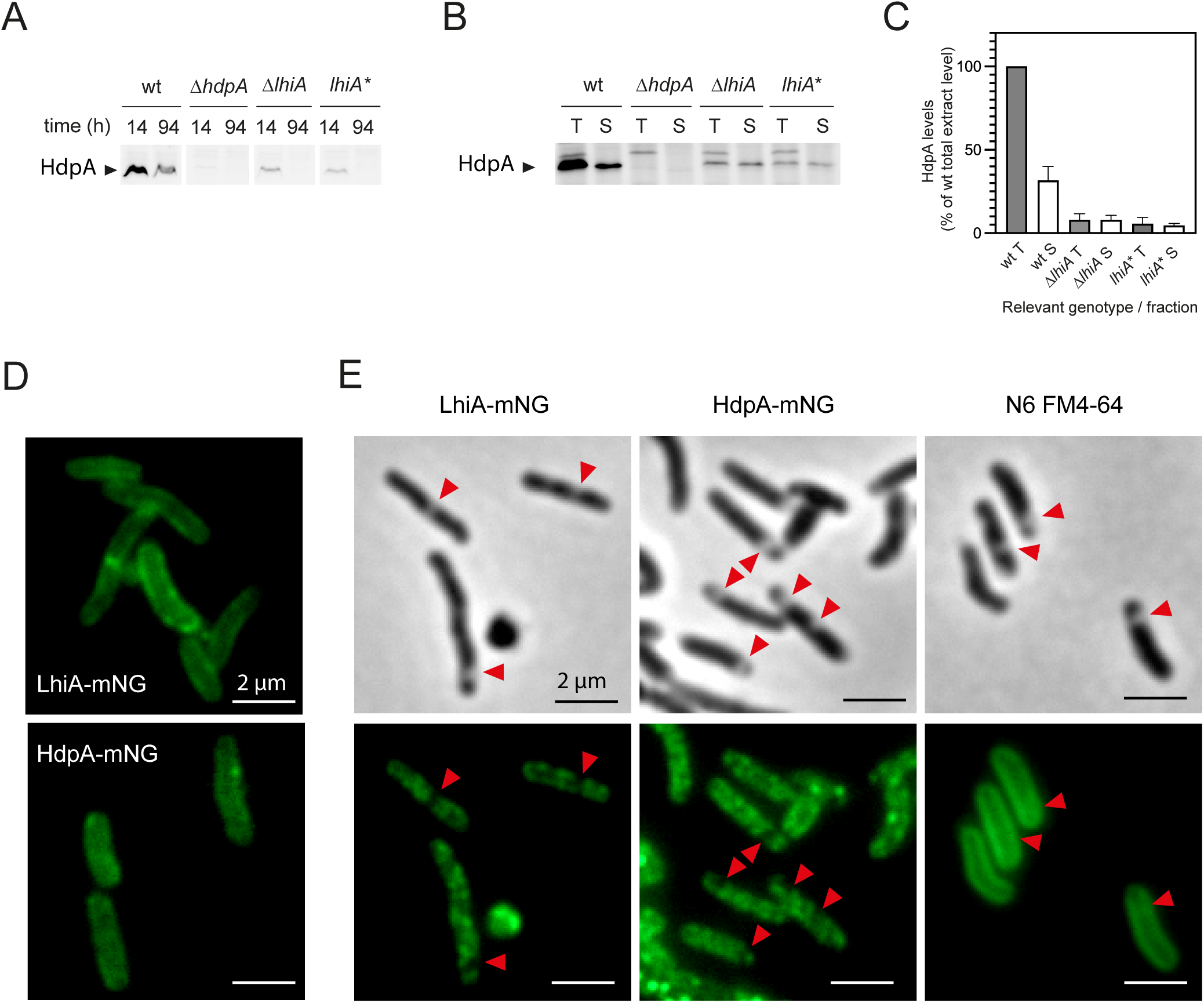
LhiA protects HdpA from proteolysis and likely recruits HdpA to the cytoplasmic membrane. A. Western blot of a gel run with total extracts of N6, N6Δ*hdpA*::*aphA3*, OMD1 and OMD2 at 14h (OD 0.56-1.11) and 94 h growth normalized by OD and probed with anti-HdpA antibodies. Ponceau staining of the membrane (loading control) is presented in Fig. S10A. B. Western blot of a gel run with total (T) and soluble (S) extracts of N6, N6Δ*hdpA*::*aphA3*, OMD1 and OMD2 at early exponential phase (OD 0.2-0.3) normalized by protein levels and probed with anti-HdpA antibodies. C. Quantification of the experiment shown in B (performed on 3 independent cultures). D. Epifluorescence microscopy images of live cells of strains OMD6 (HdpA-mNG) and OMD7 (LhiA-mNG). E. Epifluorescence and phase contrast images of plasmolyzed fixed cells of OMD6, OMD7 and FM4-64 stained N6. Red arrows show regions of invagination of the cytoplasmic membrane.

To confirm that the phenotypes observed for OMD1 are due to the deletion of the *lhiA* gene, we constructed OMD5, a derivative of OMD1 in which a wt *lhiA* allele was inserted under the control of the *ureA* promoter at an ectopic location on the chromosome, between genes *hp1499* and *hp1500*. We analyzed two phenotypes: HdpA levels at different time points and coccoid formation in strains N6, OMD1 and OMD5. The ectopic *lhiA* allele restored the HdpA levels as well as coccoid formation (Fig. S7).

### LhiA is involved in HdpA localization

PGN remodelling proteins and their regulators can be periplasmic, anchored to the outer leaflet of the cytoplasmic membrane or the inner leaflet of the outer membrane, they can also have a differential localization along the cell longitudinal axis. The reason for their localization in a particular compartment is still unclear in the majority of cases, however there is little doubt that it plays an important role in their activity, as has been shown in the case of LpoA and LpoB (25, 26). Enzyme localization can also be rendered dynamic by interaction with scaffolding proteins as shown by the example of the NlpI outer membrane lipoprotein network (27). All these examples highlight the importance of collecting more information on the subcellular localization of PGN remodeling complexes.

HdpA is expressed with an N-terminal extension which is predicted to be either a signal peptide or a transmembrane segment while LhiA is predicted to be an outer membrane lipoprotein. However, the atypical lipid content (28) as well as notable deviations in envelope biogenesis in *H. pylori* (29) warrant caution in interpreting these predictions. We therefore set out to determine where these proteins localize along the cell and to which compartment of the envelope. For that purpose, we constructed two strains, OMD6 and OMD7, in which the *hdpA* and the *lhiA* chromosomal genes were respectively fused to the *mNG* gene encoding the mNeongreen (mNG) fluorescent protein. Equal amounts of total protein extracts of these strains at early exponential phase analyzed by SDS-PAGE showed only one fluorescent band corresponding to the fusion protein in each case (Fig S8A). The similar fluorescence intensities of the two fusions suggest that in the wt context, LhiA is produced in slightly higher amounts than HdpA in exponential phase. Upon epifluorescence microscopy analysis, cells of both strains displayed a patchy peripheral labelling, suggesting that the two proteins are associated to the envelope (Fig. 5D). Accordingly, distribution of the fluorescence foci along the cell transversal axis followed a camel’s back curve with two maxima approximately one cell diameter apart (Fig. S8B). Analyzing the density of the foci in > 1000 cells showed that HdpA is present along the sidewall but excluded from the pole tip, and that it might also be the case for LhiA (Fig. S8C). To determine the compartment localization of LhiA and HdpA, we resorted to the plasmolysis technique (26, 30) in which osmotic shock is used to force inner membrane retraction from the more rigid outer membrane and create plasmolysis bays. In these conditions, both HdpA-mNG and LhiA-mNG followed the invagination of the inner membrane and disappeared from the outline of the cell lining plasmolysis bays (Fig. 5E). As a control, the lipophilic dye FM 4-64 appeared to stain the outline of the cells irrespective of the presence of plasmolysis bays, as observed before (30). Altogether, these results strongly suggest that both HdpA and LhiA localize to the inner membrane. Interestingly, the residue in position +2 of the mature LhiA protein is a lysine and not an aspartate. This confirms the notion that the rules governing lipoprotein sorting in *E. coli* cannot be extrapolated to other bacteria (31).

The localization of HdpA could be the result, either of its interaction with LhiA or of its N-terminal extension behaving as an uncleaved transmembrane segment. To distinguish between these hypotheses, we used a simple fractionation approach followed by immunodetection of HdpA in a wt, Δ*lhiA* and *lhiA\** context. Strains N6, OMD1 and OMD2 were grown to early exponential phase, cells were disrupted and soluble extracts were prepared by centrifugation and compared to total extracts. While around only 30% of HdpA was found in the soluble fraction in the wt context, the residual HdpA present in the Δ*lhiA* and *lhiA\** context was close to fully soluble (Fig. 5B, C). Along with the localisation results, this suggests that HdpA is a periplasmic soluble protein that is sequestered to the sidewall inner membrane by its interaction with LhiA.

### HdpA stays in complex with LhiA along growth

Coccoids appearing upon prolonged growth are supposed to be generated through a rupture of the sacculus (13), and HdpA could participate through its peptidase activities to this phenomenon. We considered two possible models for the role of HdpA : (i) either free HdpA in equilibrium with the LhiA-HdpA complex contributes to a progressive fragilization of the PGN along growth so that eventually the hydrolysis of one peptide bond creates a hole in the PGN sufficiently large to allow blebbing of the membranes outside of the sacculus; or (ii) a burst in HdpA enzymatic activities triggers the transition. Such a burst should happen at the start of the transition to coccoids and could either involve an increase in the concentration of the LhiA-HdpA complex, and therefore of free HdpA, or a release of free HdpA from the complex triggered by an unknown mechanism.

We therefore set out to determine whether we could observe the LhiA-HdpA complex in vivo and how it evolved during prolonged growth. For that purpose, we constructed strain OMD3, in which LhiA was produced as a C-terminus 3xFlag-tagged version. Introduction of this tag did not affect growth (Fig. S4) or the HdpA-stabilizing effect (Fig. S9) importantly. It moderately reduced the final proportion of coccoids without significantly changing the kinetics of the transition (compare Fig. S1A and the coccoid proportion curve in Fig. 6B), suggesting that the interaction with HdpA is only marginally affected. We grew strain OMD3 and N6 as a control, solublilized the proteins in detergent and immunocaptured the LhiA-Flag protein on anti-Flag magnetic beads at different time points along growth. The elution from the beads at the 24h time point was concentrated and first analyzed by SDS-PAGE and Coomassie blue staining, showing only one detectable difference between the elution of the OMD3 and that of the N6 strain, a band running at the apparent MW expected for HdpA (even the LhiA-Flag band was not detected, probably because low MW proteins diffuse more in SDS-PAGE) (Fig. 6A, left panel). This suggested that HdpA is LhiA main partner. We then compared the elutions of N6 and OMD3 to the corresponding solubilized extracts by Western blot at different time points (Fig. 6A, right panel). HdpA indeed copurified with LhiA along the growth curve, which shows that the complex exists in vivo throughout growth. No increase in the amount of the complex was observed: the amount of immunoprecipitated HdpA constantly decreased with growth as did HdpA in the solubilized extract.

**Figure 6:**
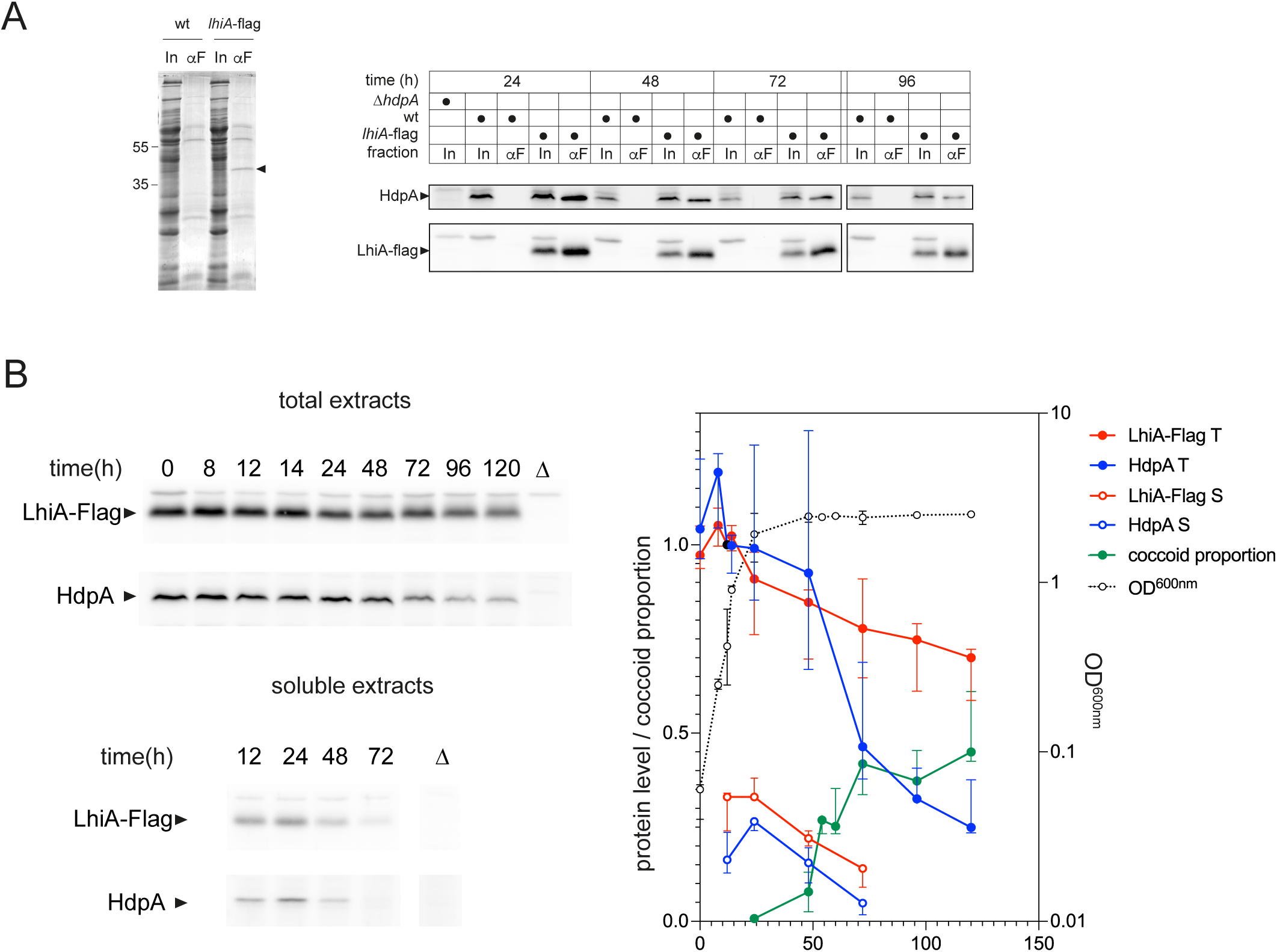
HdpA is present in vivo mainly as a membrane anchored LhiA-HdpA complex which wanes during post-exponential phase growth. Cultures of strain N6, OMD3 and N6 Δ*hdpA*::*aphA3* were grown for up to 120 h and samples were withdrawn at the indicated times. A. Anti-flag co-immunoprecipitation of extracts solubilized in 0.1% Nonidet P40 substitute. Solubilized extracts (650 µg total proteins) were adsorbed on anti-Flag magnetic beads which were washed before a pH 3 elution. Samples were run on a 13% polyacrylamide/bisacrylamide gel, and either stained with Coomassie blue (left panel, with the arrow indicating the band specific of OMD3 and relevant marker positions indicated) or transferred onto a nitrocellulose membrane and probed by anti-Flag and anti-HdpA antibodies (right panel). Relevant genotypes are indicated. In : input ; αF : elution (overloaded by a factor 2.7 for the western blot, 51 for the coomassie stained gel). B. Evolution of LhiA-Flag and HdpA protein levels along growth. Total extracts, soluble extracts and live cells of three to six independent cultures were prepared at indicated time points to analyse protein levels by western blot and coccoid proportion by phase contrast microscopy. Left panel : representative western blot of total and soluble protein extracts probed by anti-HdpA and anti-flag antibodies. Right panel : quantification (as median values with 95% confidence intervals) of the western blots and of the coccoid proportion along the growth curve. T : total protein levels. S : soluble protein levels. Protein levels are normalized with respect to the 12h total protein level (black solid circle). Ponceau staining of the membranes (loading control) is presented in Fig. S10B, C and typical microscopy images used for the determination of the coccoid proportion are shown in Fig. S11.

### LhiA and HdpA total levels decrease along growth

The analysis of the LhiA-HdpA complex in vivo is in favor of the progressive weakening hypothesis. To test the possibility of a release of free HdpA prior to coccoid transition, we followed in parallel the total and soluble contents in HdpA and in LhiA-Flag by Western blotting, while monitoring coccoid formation by phase contrast microscopy (Fig. 6B and Fig. S11). After a relative stability during exponential phase and the beginning of stationary phase, HdpA total levels decreased to reach around 25% of its maximum level at 120h, while LhiA levels regularly decreased from the exponential phase on to reach 65% of the maximum at 120h. The decrease in HdpA started with the appearance of the first coccoids, at 48h growth and lasted as the coccoid proportion increased. Similarly, soluble levels of both proteins decreased after 24h. Note that LhiA and HdpA were released in the soluble fraction in similar proportions (with respect to the total proteins), raising the possibility that the two proteins might be associated even in the soluble fraction. In conclusion, there does not seem to be any major peak of free HdpA concomitant with coccoid formation. The model that explains better our data is that LhiA-HdpA constantly release small amounts of soluble HdpA which act on the peptidoglycan substrate before being proteolysed. This activity, albeit decreasing along growth, is sufficient to progressively weaken peptidoglycan, eventually leading to the coccoid transition.

## DISCUSSION

### Role of the LhiA-HdpA complex in *H. pylori* morphogenesis

In *H. pylori*, only two enzymes are known to have the D,D-endopeptidase activity necessary to hydrolyze 4-3 cross-links generated by PGN synthases to make space for the insertion of new PGN strands: HdpA and Csd1.

Csd1 is known to act within a complex comprising its inactive homolog Csd2 and membrane protein Csd7. The effect of Csd2 and Csd7 on Csd1 activity is not known but both proteins protect Csd1 from proteolysis (32). In positive curvature regions of the envelope, there seems to be a functional coupling between the Csd1-Csd2-Csd7 endopeptidase complex and a synthetizing complex containing CcmA, membrane protein Csd5 and precursor synthesis enzyme MurF. This coupling relies on the ability of CcmA bactofilin domain to toggle between binding to Csd7 (which releases Csd1 leading to its degradation) and binding to Csd5 which anchors it to regions of positive curvature (33). Along with the cell-shape defect of *csd1* mutants and the conserved genetic association between *csd1*, *csd2* and *ccmA*, this lead to the hypothesis that Csd1 is the endopeptidase associated with a "shapesome" PGN synthesizing complex guided by CcmA filaments and localized in regions of positive gaussian curvature. Here, we show that the second endopeptidase of *H. pylori*, HdpA/Csd3, is tightly associated with a lipoprotein chaperone, LhiA, which anchors it in the cytoplasmic membrane while inhibiting its activity. As is the case for Csd1, release of HdpA from its membrane partner leads to quick degradation, underscoring the fact that endopeptidase activity can be harmful and has to be tightly regulated. Along with the fact that LhiA seems to be present in slightly higher amounts than HdpA, this indicates that the main HdpA species present in the periplasm is the LhiA-HdpA complex anchored in the inner membrane. What is the rationale for the existence of such a system? An enzyme tethered to a cytoplasmic membrane lipoprotein has one essential advantage with respect to a transmembrane enzyme: flexibility. Indeed, lipoproteins are thought to be more mobile in the membrane than transmembrane proteins, even if it has not been proven experimentally (34). The fact that the N-terminal ∼20 residue stretch of mature LhiA is not visible in the electron density map suggests that it is a disordered linker allowing HdpA to reach the PGN and/or interactants in different orientations and at different distances from the cytoplasmic membrane. This sharply contrasts with the Csd1 and Csd2 proteins, which bear N-terminal extensions predicted to be long α-helices forming a coiled-coil (see AF-O26068-F1 and AF-O26069-F1 at the Alphafold Protein Structure Database), and which are anchored by polytopic integral membrane protein Csd7, suggesting a more rigid structure. Furthermore, HdpA is not itself anchored in the membrane but only bound to a membrane anchor, providing the possibility of a release of the peptidase in the periplasm. Release events could be simply the result of law mass action applied to the LhiA-HdpA equilibrium, or they might be triggered by the presence of an appropriate substrate or by an unknown cue. However they are likely controlled in time and space by free HdpA proteolysis. Since the most striking phenotype of *lhiA* and *hdpA* mutations in exponential phase consists in the growth of aberrant poles, and given their localization in foci relatively evenly distributed along lateral PGN, our working hypothesis is that LhiA-HdpA behaves as a "task force" enzymatic complex that can be recruited anywhere along lateral PGN to rectify incorrect PGN growth. One substrate of LhiA-HdpA could be small patches of inert PGN that have been recognized as starting points for branching (35). In this line of thought, the slight excess of HdpA foci at the vicinity of the poles (Fig. S8C) might reflect a preference for inert PGN (which has to be balanced by other factors to explain exclusion from the pole tip). Whether the LhiA-HdpA complex functions autonomously, like the Lpo-class A PBP system which seems to perform a damage-repair task with a need-based localization (36), or in association with the Rod complex, as suggested by the effect of *hdpA* mutations on cell width (37) and bulgecin sensitivity (38), remains to be determined. Noteworthy, the fact that *lhiA* mutations do not recapitulate the cell width phenotype has to be taken with caution because HdpA is still produced in early exponential phase in *lhiA* defective contexts, probably dampening early phenotypes.

In addition to its role in cell shape maintenance, HdpA is the only PGN endopeptidase known to play a role in the transition to the coccoid form. Here, we show that LhiA plays a positive role in this transition through its interface with HdpA, i.e. most likely through its stabilization of HdpA. We also show that the transition to coccoids happens by progressive weakening of the sacculus probably due to PGN synthesis being insufficient to compensate for PGN hydrolysis stemming from low-key endopeptidase activity of free HdpA released by LhiA-HdpA. We view the LhiA-HdpA complex as a way to store endopeptidase activity generated in excess while nutrients are available so that it can continue to be used for PGN remodeling in a phase where energy resources are limited, without the need for protein synthesis and translocation. Strikingly, inhibition of cell wall synthesis by different compounds in *V. cholerae* results in endopeptidase-dependent formation of cells that resemble the *H. pylori* coccoid form (16), and one of the endopeptidases triggering this transition is HdpA homolog ShyA. Note also that the problem of balancing peptidoglycan synthesis and hydrolysis during stationary phase seems to be solved in two different ways according to the organism: either by reducing the surface of peptidoglycan by dwarfism and a conversion to cocci with an intact but reduced peptidoglycan surface, or by keeping the same surface of sacculus but eventually turning into a "peptidoglycan-naked" coccoid form as in *H. pylori*.

Finally we observe that proteolysis combined with scaffolding protein interaction seems to be a common theme in PGN endopeptidases. LhiA-HdpA system shares similarities with the control of the MepS peptidase by the NlpI outer membrane lipoprotein and the Prc protease in *E. coli*, one of the rare other endopeptidase complexes for which a crystal structure is available (39). One convergence point between these systems is that the effect of the scaffolding protein on enzymatic activity (positive in the case of NlpI-MepS, negative in the case of LhiA-HdpA), is opposite with respect to its effect on protein stability (negative in the case of NlpI-MepS, positive in the case of LhiA-HdpA) (27, 40). This equilibrium between opposite effects at the protein level might represent an efficient way to fine-tune the activity of the enzyme in response to environmental or periplasmic cues, with a better reactivity compared to transcription regulation (40). The opposite roles of the scaffolding proteins with respect to proteolysis as well as the difference in localization might reflect functional differences between the HdpA and MepS peptidases.

### A new peptidoglycan remodeling complex in Campylobacterota

Campylobacterota are an extremely diverse phylum of bacteria found in environments as different as marine hydrothermal vents and animal hosts, the latter exemplified by *Helicobacter* and *Campylobacter* genera. Several of them have been found to undergo a transition from a (helical) rod to a coccoid form and HdpA is well conserved in the phylum(12). Search for LhiA orthologs is complicated by the fact that the thioredoxin fold is extremely widespread (41). We conjectured that LhiA-like HdpA regulators might display structural features absent in other thioredoxins, therefore we performed a Foldseek search (42) of the AFDB50 database seeded with the Alphafold model of the LhiA protein from *H. pylori* 26695. We obtained numerous hits in the Campylobacterota, including sequences from remote genera like *Sulfurovum* or *Nitratiruptor*. Among the 102 first hits, one of the most conserved motifs was the GXXPXEML sequence at positions 169-175 (LhiA^N6^ numbering, Fig. S12). In the crystal structure of the complex, G169 and P172 most likely favor the formation of the β7α4 hairpin which builds up the main interaction surface of LhiA with HdpA, while E174 and M175 are key residues in the interaction (see above), and L176 contributes to the packing of the β7α4 motif against the rest of the LhiA structure, suggesting that the GXXPXEML motif is a marker of bona fide HdpA-regulating LhiA orthologs. Indeed, Alphafold 2 prompted with LhiA:HdpA sequences from remote bacteria such as Nitratiruptor (22% identity for LhiA, 45% for HdpA) generated models very similar to the crystal structure of the *H. pylori* HdpA-LhiA complex (Fig. S13). It seems therefore likely that members of the LhiA family are HdpA regulators in a large part of Campylobacterota. Note that the catalytic cysteins are conserved in all Campylobacterota with the exception of the group of gastric *Helicobacter* (Fig. S14), suggesting a complete repurposing of LhiA from a thioredoxin to a PGN remodeling activity in the latter, while the two activities might still be linked in the rest of Campylobacterota.

### Mechanism of activation of the ShyA/MepM/HdpA endopeptidases

Although peptidoglycan is a major target of antibiotic design, the search for antimicrobials has been largely biased towards compounds inhibiting penicillin-binding proteins. One alternative or complementary approach could be the search for drugs that would increase the activity of hydrolases, which underscores the importance of understanding their mechanism of activation. In the case of M23 peptidases of the ShyA/MepM/HdpA family, activation occurs through an opening of the structure resulting in the release of auto-inhibitory contacts between domain 1 and catalytic domain 3 (10). However the early events that trigger this opening are unknown. Assuming that the 3 globular domains of these proteins behave as rigid bodies with 2 hinges corresponding to the 1-2 and 2-3 linkers, two mechanisms can be considered for the first step of activation: a deformation of hinge 1-2 or of hinge 2-3. The fact that contacts between domain 1 and 2 (Fig S15A) are less numerous than contacts between domain 2 and 3 (Fig S15B) suggests that it is less energetically costly to change the relative orientations of domains 1 and 2 than domains 2 and 3. Note also that the polypeptide chain exiting domain 3 comes back to cap domain 2, which is expected to strengthen the interaction between domain 2 and 3. Our work supports the idea that endopeptidase activation starts with a deformation of hinge 1-2 since (i) LhiA strongly inhibits HdpA in a purified system and (ii) its location in the crystal structure is expected to prevent deformation of the 1-2 hinge, but not the 2-3 hinge (Fig S15C). In conclusion, we propose that the first event in the relief of auto-inhibition of M23 peptidases of the HdpA family is a movement of domain 1 away from a rigid structure made of domain 2 and 3, and that preventing this movement allosterically freezes the protein in the autoinhibited conformation. Conversely, a drug designed to interact with domain 1 or 2 in the same region as LhiA and to destabilize the 1-2 contacts is expected to trigger activation of these peptidases.

## MATERIALS AND METHODS

### Strain and plasmids

Strain DV900(DE3) (22) was reconstructed from DV900 (43) by transduction using a λDE3 lysogenization kit (Novagen). *H. pylori* strains were cultured at 37°C in a medium containing 90% Brain-Heart infusion (BHI, Oxoid) and 10% decomplemented fetal calf serum (FCS, Eurobio) (BHI/FCS) under an atmosphere containing 6% O_2_ and13% CO_2_.

*H. pylori* strains (Table 1) were constructed by standard techniques, using megaprimer PCR or Gibson assembly to generate DNA fragments containing the desired mutation along with cassettes containing genes *aphA*-3 or *aac(3)IV* (see Table S3) These DNA fragments were used for natural transformation of the relevant *H. pylori* strain. The correct transformants were selected for kanamycin or gentamicin resistance, respectively, and final constructs were checked by sequencing.

**Table 1.**

| Strain | relevant genotype | Reference |
| --- | --- | --- |
| DV900(DE3) | $\Delta_{ponB\ dacA\ dacB, dacC\ dacD\ pbpG\ ampH\ ampC\ \Delta_{pbp4B}$ (λDE3) | (43) and this work |
| N6 | wt *H. pylori* strain | (53) |
| N6hp0506ΩKm | N6$\Delta_{hdpA::aphA3}$ | (17) |
| OMD1 | N6 $\Delta_{lhiA::aphA3}$ | This work |
| OMD2 | N6 *lhiA* * *aphA3* | This work |
| OMD3 | N6 *lhiA-flag3 aphA3* | This work |
| OMD5 | OMD1 $\Omega_{(ureAp-lhiA\ aac(3)IV)}$ | This work |
| OMD6 | N6 *hdpA-mNeongreen aphA3* | This work |
| OMD7 | N6 *lhiA-mNeongreen aphA3* | This work |

Plasmids pOD1, pOD2, pOD2 *E174R M175R* and pOD3 derivatives of pET24b(+) were introduced in the relevant DE3 prophage harbouring strain and used for production of HdpA^S^-LEHHHHHH, LhiA^S^-SAHHHHHH, LhiA^S^*-SAHHHHHH and fusion protein LhiA^S^-HdpA^S^-LEHHHHHH, respectively. pGEX*hdpA* is a pGEX-4T-1 derivative used for production of GST-HdpA in the BLi5 strain (Novagen). HdpA^S^ and LhiA^S^ designate the polypeptides starting with a methionine followed by positions 29-406 of HdpA and positions 25-185 of LhiA, respectively (sequences from strain 26695 (44)). Plasmid constructs are presented in Table S4.

### Protein purification

Tagged proteins were purified by standard affinity chromatography (using His-Trap columns for His-tagged proteins or Glutathion-Sepharose™ 4B for GST fusions) and exclusion chromatography procedures. For the purification of His-tagged HdpA^S^ and LhiA^S^ used in the enzymology and exclusion chromatography experiments, we used expression strain DV900(DE3) to avoid contamination by PBP endopeptidase or carboxypeptidase activities. Indeed, LhiA^S^ showed no peptidase activity, while all peptidases activities of the newly purified HdpA^S^ were abolished by EDTA (Fig. S2). For all other purifications of His-tagged proteins, BL21(DE3) was used as a host.

Protein concentrations were determined by measuring the absorbance at 280 nm of dilutions in 6M guanidinium hydrochloride using their theoretical extinction coefficient.

The HdpA^S^-LhiA^S^ complex was formed by mixing stoichiometric amounts of the two His-tagged proteins, concentrating on a Vivaspin 6 (Cytiva) and further purified on a Superdex 200 10/30 GL 24 mL column equilibrated with a 7 mM Hepes-KOH, pH8.0 buffer containing 10% glycerol and 0.35 M NaCl. The complex was concentrated again to 38 mg/mL and used for crystallisation trials after adding 0.5 mM Tris(2-carboxyethyl)phosphine hydrochloride (TCEP).

For HdpA^S^-LhiA^S^ fusion protein purification, an ion exchange step was added between the affinity and the exclusion chromatography.

GST and GST-HdpA were purified by standard procedures from cultures of BLi5 (pGEX-*hdpA^S^*) and BLi5 (pGEX) (17), respectively.

### Pull-down assays

GST-HdpA^S^ or GST were allowed to bind Glutathion-Sepharose™ 4B resin equilibrated in buffer A (see supplementary Materials and Methods for the composition of the buffers) + 1 mM DTT at a concentration of 2 mg (29 nmol) per mL of resin (GST-HdpA^S^, or 1.8 mg (65 µmol) per mL of resin (GST) O.N. at 4°C. GST- or GST-HdpA^S^-carrying slurry was divided in three 150 µL columns which were washed with Buffer A, then with buffer D + 1 mM DTT. His-tagged LhiA^S^, LhiA^S^* (4.3 nmol in 150 µL, buffer D + 1 mM DTT) or 150 µL buffer D + 1 mM DTT were loaded on the 3 columns, which were washed with 3 column volumes of buffer D + 1 mM DTT and eluted with 3 column volumes of buffer D + 1 mM DTT + 10 mM reduced glutathion.

### Analytical exclusion chromatography

Proteins were incubated for 10 min at room temperature in a 43 mM Tris-HCl buffer containing 4.5 mM Hepes-NaOH, 10 % glycerol, 490 mM NaCl and injected on a Superdex 200 PC3.2/30 column (GE Healthcare) equilibrated in buffer A mounted on an Ettan LC system (GE Healthcare) run at room temperature and at a 40 µl/min flow-rate. 50 µL fractions were collected. Apparent molecular weights were deduced from a standard curve obtained by running β-amylase (200 kDa), bovine serum albumin (66kDa) and cytochrome c (12.5 kDa) on the column in the same conditions.

### Peptidase assays

HPLC-purified anhydromuropeptides at different concentrations were incubated with the relevant proteins in a buffer containing 25 mM HEPES pH8.0, 5 mM NaCl and 5% glycerol. Initial reaction velocities were measured at least in triplicate per substrate concentration. Enzyme concentration was 0.4 µM. Reactions were stopped by adding trifluoroacetic acid (TFA) at a final concentration of 0.2%. Products of the reaction were separated by reverse phase HPLC using a a Hypersil C18 Gold column (250 x 4.6 mm, 5-µm particle size, Thermo Fisher Scientific) mounted on a Shimadzu LC-20 system. The separation conditions used were a 0-25% acetonitrile gradient in 0.05% TFA over 135 min at 52 °C and a flow rate of 0.5 ml/min. Quantification of the substrate and products was done by integration of the absorbance peaks at 206 nm, and all anhydromuropeptides identifications were validated by MS/MS. The ratio of the peak areas was used to calculate the proportion of consumed substrate. Molar extinction coefficients of anhGM5 and anhGM4-anhGM4 used to determine the substrate concentrations were estimated to 12400 and 24000 M^-1^cm^-1^, respectively, by measuring the absorbance and the concentration of a stock solution in H_2_O. Absorbance was measured at 206 nm on a Nanodrop™ (Ozyme). Concentration was estimated by 1H-NMR spectroscopy, by adding D_2_O to 5%, an internal standard, trimethylsilylpropanesulfonate (DSS) to 1 mM, and comparing the integral of the amide proton of anhydro-MurNAc at 7.4 ppm with the integral of the methyl protons of DSS at 0 ppm. Concentrations of anhGM4-anhGM5 and anhGM5 were deduced from their absorbances at 206 nm using the molar extinction coefficients corrected as shown in (45).

### Crystallization and diffraction data collection

Crystallization condition screenings and hits optimization were performed at the Crystallography Core Facility of the Institut Pasteur (46). Initial crystallization screening trials were carried out by the sitting drop method with a Mosquito (TTP Labtech) automated nanoliter dispensing system. Crystals were grown using the vapour diffusion method at 18°C, mixing equal volumes of protein and reservoir solutions. Initial crystallization screening trials were carried out by the sitting drop method with a Mosquito (TTP Labtech) automated nanoliter dispensing system. After manual optimization of the obtained initial hits, single crystals corresponding to the LhiA^S^ HdpA^S^ complex and to the LhiA^S^-HdpA^S^ fusion grew in 100 mM Tris-HCl (pH7.5), 3.0 M Na formate and 100 mM Tris-HCl (pH 8.5), 30% w/v PEG 10000 respectively. Crystals were flash-frozen using the crystallization solution supplemented with 33% (v/v) of ethylene glycol as a cryoprotectant and then stored in liquid nitrogen until data collection. X-ray diffraction data were collected at beamlines PROXIMA 1 and PROXIMA 2a (Synchrotron SOLEIL, St. Aubin, France) and processed with autoPROC (47).

### Structure determination and refinement

The crystal structure of LhiA^S^ HdpA^S^ complex and LhiA^S^-HdpA^S^ fusion were solved by molecular replacement with Phaser (48) using AlphaFold2 (49) models as search templates. The models were refined through several interactive cycles using Coot (50) and Buster-TNT (51). Data collection and refinement statistics are summarized in Table S1 and S2. Atomic model coordinates and structure factors have been deposited in the Protein Data Bank under the accession codes 9RBX (LhiA^S^ HdpA^S^ complex) and 9RBY (LhiA^S^-HdpA^S^ fusion). Figures showing the crystallographic models were generated with Pymol (Schrödinger, LLC).

### Microscopy

Bacteria from cultures in BHI/FCS were concentrated by centrifugation, and either recovered in BHI or fixed in 70% ethanol for 15 min at -20°C, pelleted and recovered in TBS (Tris buffered saline). For plasmolysis experiments, cells were recovered in 30 µL TBS and within 1 min fixed with 70 µL 100% ethanol at -20°C for 15 min, collected and recovered in TBS or TBS containing 10 µg/mL FM4-64 (Thermofisher Scientific). For analysis, cells were spotted on a 1% agar pad in PBS (Phosphate-buffered saline). Phase contrast or epifluorescence images were acquired using an inverted microscope Axio Observer (Zeiss), equipped with a 100x/1.40 oil immersion objective and a Colibri LED excitation source. All fluorescence images were acquired using Filter set 61 HE (Zeiss). Images were analyzed using MicrobeJ (52) as indicated in Supplementary Material.

### Analysis of total and soluble HdpA and LhiA

*H. pylori* strains were inoculated at OD 0.05 in BHI/FCS medium and grown at 37°C in 6% O_2_ under agitation. At various times 1 OD of cells was harvested by centrifugation at 9000 x g for 10 min at 20°C. Pellets were recovered in a 50 mM Tris-HCl buffer (pH8.0) containing 10% glycerol, 200 mM NaCl and Complete w/o EDTA x2 (buffer E), sonicated, and protein content was analyzed by BCA assay (BioRad). Samples were either directly analyzed after normalization for protein content, by SDS-PAGE on a 13% acrylamide/bisacrylamide gel (total extracts only), or centrifuged for 2h at 21000 x g, and equal volumes of supernatant and total extract were analyzed by SDS-PAGE after normalization for total extract protein content.

Proteins were transferred on nitrocellulose membranes (Bio-Rad) which were stained with Ponceau Red, destained and blocked overnight at 4°C in 10 mM Tris-HCl-pH8.0, 150 mM NaCl, Tween-20 0.1%. Membranes were probed with Guinea pig-raised anti-HdpA polyclonal antibodies or with anti-Flag**^®^** M2 monoclonal antibody (F3165, Sigma-Aldrich). Secondary antibodies were HRP conjugates and protein bands were analyzed on a ChemiDoc**™** (Bio-Rad) using chemiluminescent substrate ECL femto (Pierce).

### Immunoprecipitation

*H. pylori* strains were grown as for total HdpA/LhiA analysis, but 4 OD of cells were collected at different times and pellets were recovered in buffer E supplemented with 0.1% Nonidet P40 substitute (AMRESCO) and disrupted by sonication. Protein concentration was determined by BCA assay. Extracts were centrifuged for 2h at 21000 x g. Volumes of supernatant corresponding to 650 µg of total proteins were applied to 10 µL (bead volume) anti-Flag**^®^** M2 magnetic beads (Millipore) for 2h at 4°C equilibrated in buffer. Beads were washed 4 times with 400 µL buffer E + 0.1% Nonidet P40 substitute. Bound proteins were eluted with 100 µL glycine-HCl, pH 3.0 and immediately neutralized with 10 µL 1 M Tris pH 8.0. Samples were then analyzed by SDS-PAGE (13%) either directly (for immunoblotting as above) or after trichloroacetic acid precipitation and acetone wash (for Coomassie Blue staining).

## ACKNOWLEDGEMENTS

We thank Juan Ayala for the gift of strain DV900. We are grateful for Kerime Özkan for technical assistance. We acknowledge SOLEIL for provision of synchrotron radiation facilities, and we thank the staff of beamlines PROXIMA-1 and PROXIMA-2A for assistance. The authors are grateful to the Staff of the Crystallography platform at the Institut Pasteur for robot-driven crystallization screenings. We thank Evelyne Richet for her comments on the manuscript. The Boneca laboratory was supported by the following programs: Investissement d’Avenir program, Laboratoire d’Excellence “Integrative Biology of Emerging Infectious Diseases” (ANR-10-LABX-62-IBEID); FRM Grant Programme d’Urgence (DBF20160635726); and Équipe FRM Grant (EQU202403018034).

## Data Availability Statement

Crystallographic model coordinates and structure factors were deposited in the Protein Data Bank with accession codes 9RBX (LhiA^S^ HdpA^S^ complex) and 9RBY (LhiA^S^-HdpA^S^ fusion). All other data are included in the article and in the Supplementary Information.

