## Supplementary Information for "Architecture of a peptidoglycan peptidase complex involved in morphological transition in *Helicobacter pylori*"

**This PDF file includes:**

Supporting Information text
Figures S1 to S15
Tables S1 to S6
SI References

### Supporting Information Text

#### Material and Methods

**Strains.** Constructs of deletions and insertions in the *H. pylori* N6 strain were carried out as follows: parental strains (in bold in the genotype column) were transformed with DNA from the Megaprimer PCRs or Gibson assemblies by natural transformation followed by selection on blood agar plates containing 20 µg / mL kanamycin (*aphA*-3 cassette) or 5 µg / mL gentamicin (*aac(3)IV* cassette). Primers are listed in Table S6

*Megaprimer constructs*

Amplicons obtained by the relevant L (primers L1 and L2, L genomic DNA) and R (primers R1 and R2, R genomic DNA) PCRs were used as megaprimers to amplify the indicated antibiotic resistance cassette along with primers L1 and R2 (Table S3). To avoid polar effects, the final construct was designed so that the antibiotic resistance gene (i) was transcribed in the same direction as the upstream gene (ii) either had its START codon overlapping with the STOP codon of the upstream gene or had its START codon preceded by a Shine Dalgarno sequence (1).

*Gibson constructs*

They were obtained by Gibson assembly of the indicated PCRs (Table S3).

OMD6 (N6 *hdpA-mNeongreen-aphA3*) was constructed by transforming the N6 strain with the pBR322-*HP0506-mNG-aphA3-HP0507* suicide plasmid and selecting for Kanamycin resistance. In the obtained strain *hdpA* is fused to *mNeongreen* (sequence encoded at the junction LLEGFGEED, HdpA residues underlined), and the fusion is translationnally coupled to *aphA3*, itself translationnally coupled to HP0507, as are *hdpA* and *HP0507* in the native sequence (i.e. by TA**ATG** sequences, where the STOP codon of the upstream gene is underscored and the start codon of the downstream gene is in bold).

**Plasmids.** pBR322-*HP0506-0507* was constructed by cotransformation of amplicons generated by (i) inverse PCR of pBR322 with primers pBR1 and pBR2 (ii) a PCR of N6 genomic DNA using primers pBR506 and pBR507 encompassing the *hdpA* and *HP0507* genes. pBR322-*HP0506-mNG-HP0507* was then constructed by cotransformation of amplicons generated by (i) inverse PCR of pBR322-*HP0506-0507* with primers 506R2 and 507F2 (ii) a PCR of plasmid OmpA-mNG-sfTq2ox (2) DNA using primers 506-Neon and 507-Ne-sfT. OmpA-mNG-sfTq2ox was a gift from Tanneke den Blaauwen (Addgene plasmid # 124220 ; http://n2t.net/addgene:124220; RRID:Addgene_124220). Finally, pBR322-*HP0506-mNG-aphA3-HP0507* was then constructed by cotransformation in E. coli DH5α of amplicons generated by (i) inverse PCR of pBR322-*HP0506-mNG-HP0507* with primers NeonR1 and 507F3 (ii) a PCR of pUC18K with primers Neon-Kan and Kan-507.

Other constructs (Table S4) were performed by inserting a DNA fragment obtained by PCR on *H. pylori* genomic DNA using the indicated primers introducing the indicated restriction sites at the extremities in the corresponding sites of linearized pET24b(+) (Novagen). Mutations causing the replacement of E174 and M175 of LhiA with two arginine residues in pOD-2 were obtained by inverse PCR with primers 762EMRR-F26695 and 762EMRR-Rev26695, DpnI treatment and transformation into DH5α, resulting in the replacement of the corresponding codons GAAATG bya CGTCGC sequence. The corresponding plasmid is called pOD-2 *E174R M175R*.

**Protein purification.** For the purification of His-tagged HdpA devoid of its hydrophobic segment (HdpA$^{S}$-H$_6$) used in the enzymology and exclusion chromatography experiments, DV900(DE3) (pOD-1) was grown in LB medium containing 10 µg/mL Kanamycine sulfate at 37°C to OD 0.5, induced with 0.1 mM IPTG and allowed to grow at 22°C for 24h. Cells were collected by centrifugation and recovered in a 50 mM Tris-HCl, pH8.0 buffer containing 10% glycerol and 0.5 M NaCl (buffer A) supplemented with one tablet Complete without EDTA (Roche) per 50 mL. Cells were disrupted with a French press at 16000 psi, and soluble extracts (ca 5000 OD) from a centrifugation at 144000 x g, 4°C were loaded on a 5 mL HisTrap column (GE Healthcare) equilibrated in a 25 mM Tris-HCl, pH8.0 buffer containing 10% glycerol and 0.5 M NaCl (buffer A'). The column was washed with 50 column volumes of A and eluted by steps of buffer B (Buffer A + 0.5 M imidazole). The protein eluted between 200 and 300 mM imidazole. Pooled His-Trap fractions were concentrated on a Vivaspin 50 (Cytiva, MWCO 10kDa) and further purified on a Superdex 200 10/30 GL (GE Healthcare) 24 mL column equilibrated with a 20 mM 4-(2-hydroxyethyl)-1-

piperazineethanesulfonic acid (HEPES)-KOH, pH8.0 buffer containing 10% glycerol and 0.5 M NaCl.
His-tagged LhiA devoid of its signal peptide and lipidated cysteine (LhiA$^S$-H$_6$) was purified from DV900(DE3) (pOD-2) grown in LB medium containing 25 µg/mL Kanamycine sulfate at 37°C to OD 1, induced with 1 mM IPTG and allowed to grow at 37°C for 6h. Collection and disruption of the cells, as well as His-Trap chromatography were carried out as for HdpA$^S$ except that the protein eluted between 150 and 200 mM imidazole. The polishing step by superdex 200 chromatography was also performed as for HdpA$^S$.
Protein concentrations were determined by measuring the absorbance at 280 nm of dilutions in 6M guanidinium hydrochloride using their theoretical extinction coefficient.
LhiA-HdpA hybrid protein was purified from BL21(DE3) (pOD-3) cells grown at 30°C in LB medium supplemented with 50xM, 0.5% glycerol, 2mM $MgCl_2$ and 50 µg/mL kanamycine sulfate, induced at OD 1.6 with 0.1 mM IPTG. 150 and 300 mM imidazole. Collection and disruption of the cells, as well as His-Trap chromatography were carried out as above except that 14000 OD starting material was used and the protein eluted between 150 and 300 mM imidazole. Pooled fractions were adjusted to 0.25 M NaCl with a buffer containing 20 mM Hepes-KOH and 10 % glycerol and purified by cation exchange on a 1 mL Resource S column (Cytiva). Resource S fractions were then pooled, concentrated on Vivaspin (30 kDa MWCO, Cytiva) and further purified on a Superdex 200 10/30 GL (GE Healthcare) 24 mL column equilibrated with a 20 mM HEPES-KOH, pH8.0 buffer containing 10% glycerol and 0.4 M NaCl.
Purification of HdpA$^S$, LhiA$^S$ and LhiA$^{S*}$ for all other purposes (complex crystallisation, pull-down experiments) were done as above with the following changes : proteins were purified from a BL21(DE3) host instead of DV900(DE3), transformed with pOD-1, pOD-2 or pOD-2 *E174R M175R* , respectively. Growth medium was LB medium supplemented with 50xM (3) 0.5% glycerol, 2mM $MgCl_2$ and 50 µg/mL kanamycine sulfate for both strains, induction was with 0.5 mM IPTG at OD 2.5 (HdpA) or 1 mM at OD 1.5 (LhiA) and growth was allowed to proceed for 24h at 22°C (HdpA) or 37°C (LhiA). For LhiA, buffer A and A' were replaced by buffer C (50 mM Tris-HCl, pH8.0, 10% glycerol and 0.2 M KI) and buffer B by buffer C + 0.5 M imidazole. The superdex 200 chromatography buffer was 10 mM Hepes-KOH, pH8.0, 10% glycerol 0.4 M NaCl (Buffer D).
GST and GST-HdpA were purified from strains BLi5 (pGEX-*hdpA*) and BLi5 (pGEX-4T-1), respectively. Cultures were grown in LB medium supplemented with 50xM + 0.5% glycerol + 0.5% glycerol and 2 mM $MgCl_2$ at 37°C, induced at OD 2 and put at 20°C O.N. Cells were collected by centrifugation at 144000 x g and recovered in a 50 mM Tris-HCl, pH8.0 buffer containing 10% glycerol and 0.3 M NaCl supplemented with 1 mM dithiothreitol (DTT), 0.2% Nonidet® P-40 substitute (AMRESCO) and one tablet Complete without EDTA (Roche) per 50 mL. Cells (7000-12000 OD) were disrupted with a French press at 16000 psi and centrifuged at 16000 g for 35 min. Soluble extracts were incubated with ~3 mL Glutathion-Sepharose™ 4B resin (GE Healthcare) equilibrated in buffer A for 2h30 at 4°C. Resins were washed with 40 mL buffer A + 5 mM DTT and eluted with buffer A + 5 mM DTT + 10 mM reduced glutathion. Proteins were concentrated by Vivaspin (10 MWCO) and further purified on a Superdex 200 10/30 GL (GE Healthcare) equilibrated in buffer A. GST-HdpA was reconcentrated and further purified on a Superdex 75 10/30 GL (GE Healthcare) equilibrated in buffer A.
**Microscopy image analysis.** Microscopy images were analyzed using the MicrobeJ plugin of the Fiji software (4) two sets of parameters were used to identify bacteria and distinguish between rod and coccoid shape (Table S5). Bacteria identifed were all checked by visual inspection to discard badly segmented bacteria, incorrect identification or out of focus bacteria.
For the identification of branched cells, use of the branching option of MicrobeJ identified only a minority of branched cells. Since branched cells do not represent an important proportion of the cells in most of the cases we used the parameters of Table S5 to identify and count bacteria, while branched cells (i.e. cells that have at least three poles) were counted manually. Branched cells that were not identified by MicrobeJ were added to the final total count before calculating the proportion.
For the analysis of fluorescent cells, cells were identified from the phase contrast channel using the same parameters and visual check as above. Fluorescent foci were identified as local maxima of the fluorescence channel with parameters tolerance = 500 and Z-score = 20. Distribution of the foci along the cell diameter and density of foci within the cells were analyzed using the subcellular localization tool with parameter bandwidth set to 0.05. For the analysis of curved cells, the analysis

was restricted to cells with only one curvature (C-shaped), selected by visual inspection among MicrobeJ identified bacteria and oriented using the polarity tool of MicrobeJ (using side length as a criterion).

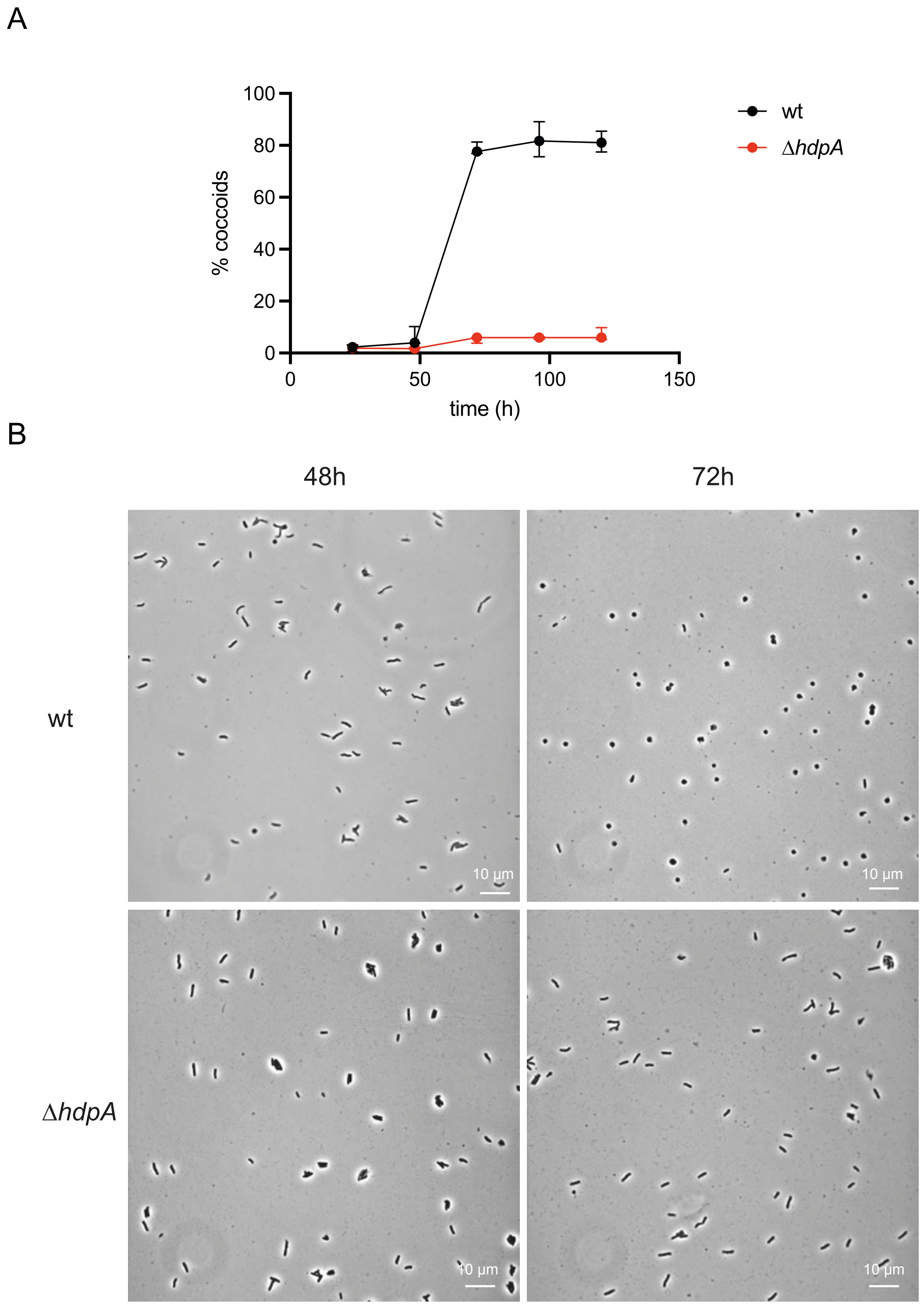


**Fig. S1.** Transition to the coccoid form in the *H. pylori* N6 (wt) and N6*hp0506*ΩKm (Δ*hdpA*) strain. Both strains were grown as described and phase contrast microscopy images of live cells were taken at the indicated times. A. Quantification of the coccoid proportion determined using ImageJ (median of 3 independent experiments with 95%CI, >150 cells counted at each time point for each experiment). B. Representative microscopy images of cells from both strains at 48h and 72h.

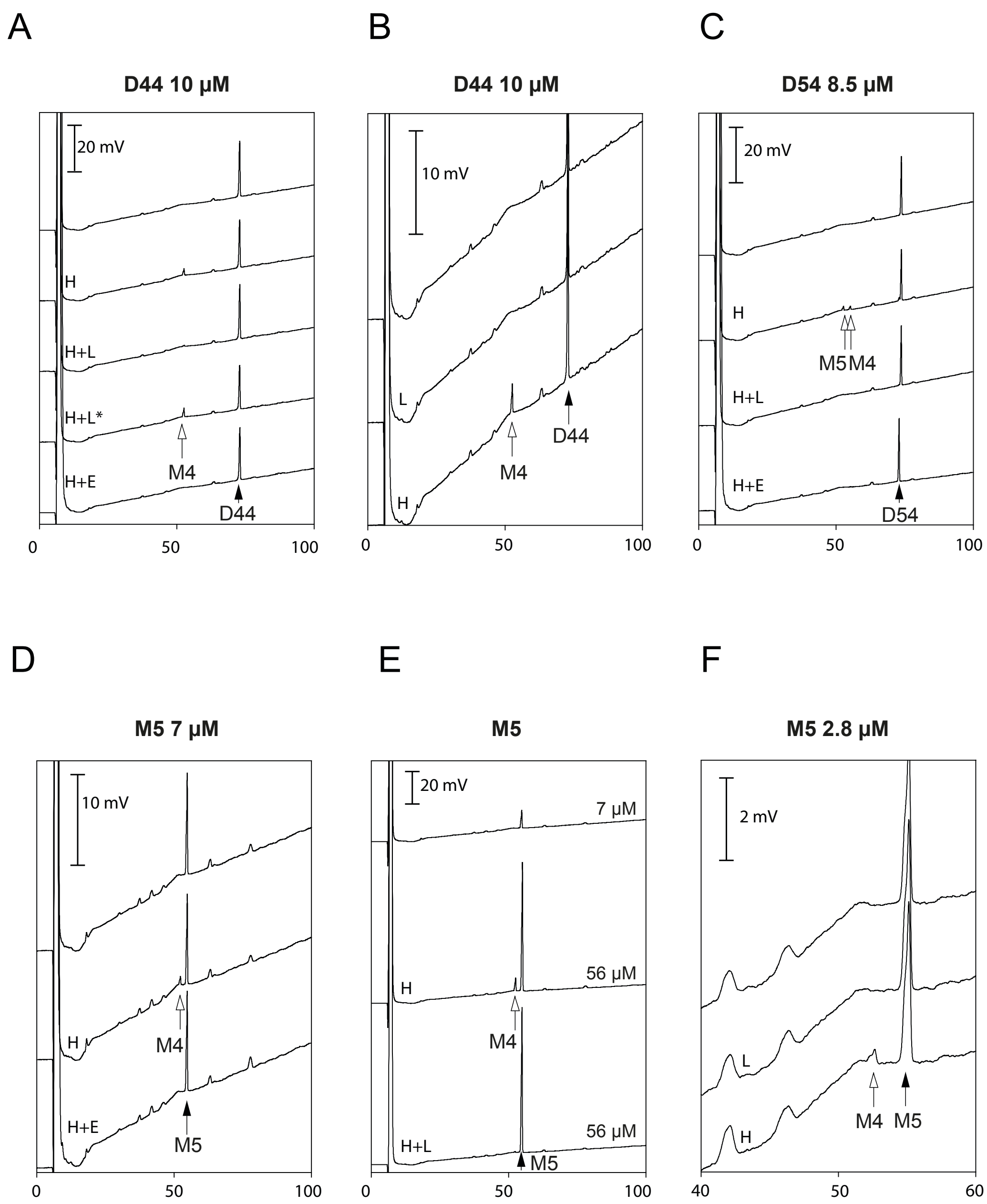


**Fig. S2.** Enzymatic activities of HdpA$^S$ (HPLC traces). Black and white arrows indicate peaks corresponding to the substrate and the product(s) of the reaction, respectively. A. Original traces used for Fig. 3A plus a trace corresponding to a control digestion of D44 by HdpA$^S$ in the presence of 20 mM EDTA (H+E). B. LhiA$^S$ is not contaminated by endopeptidase activity. D44 (10μM) was incubated alone (top trace), with 0.6 μM LhiA$^S$ (L) or 0.4μM HdpA$^S$ (H). C. Original traces used for Fig. 3B plus a trace of a control digestion of D54 by HdpA$^S$ in the presence of 20 mM EDTA (H+E) D. HdpA carboxypeptidase activity is inhibited by EDTA. M5 (2.8 μM) was incubated alone (top trace), with 0.4 μM HdpA$^S$ (H) or 0.4 μM HdpA$^S$ and 20 mM EDTA (H+E). E. Complete traces used for Fig. 3C. F. LhiA$^S$ is not contaminated by carboxypeptidase activity. M5 (2.8 μM) was incubated alone (top trace), with 0.6 μM LhiA$^S$ (L) or 0.4μM HdpA$^S$ (H).

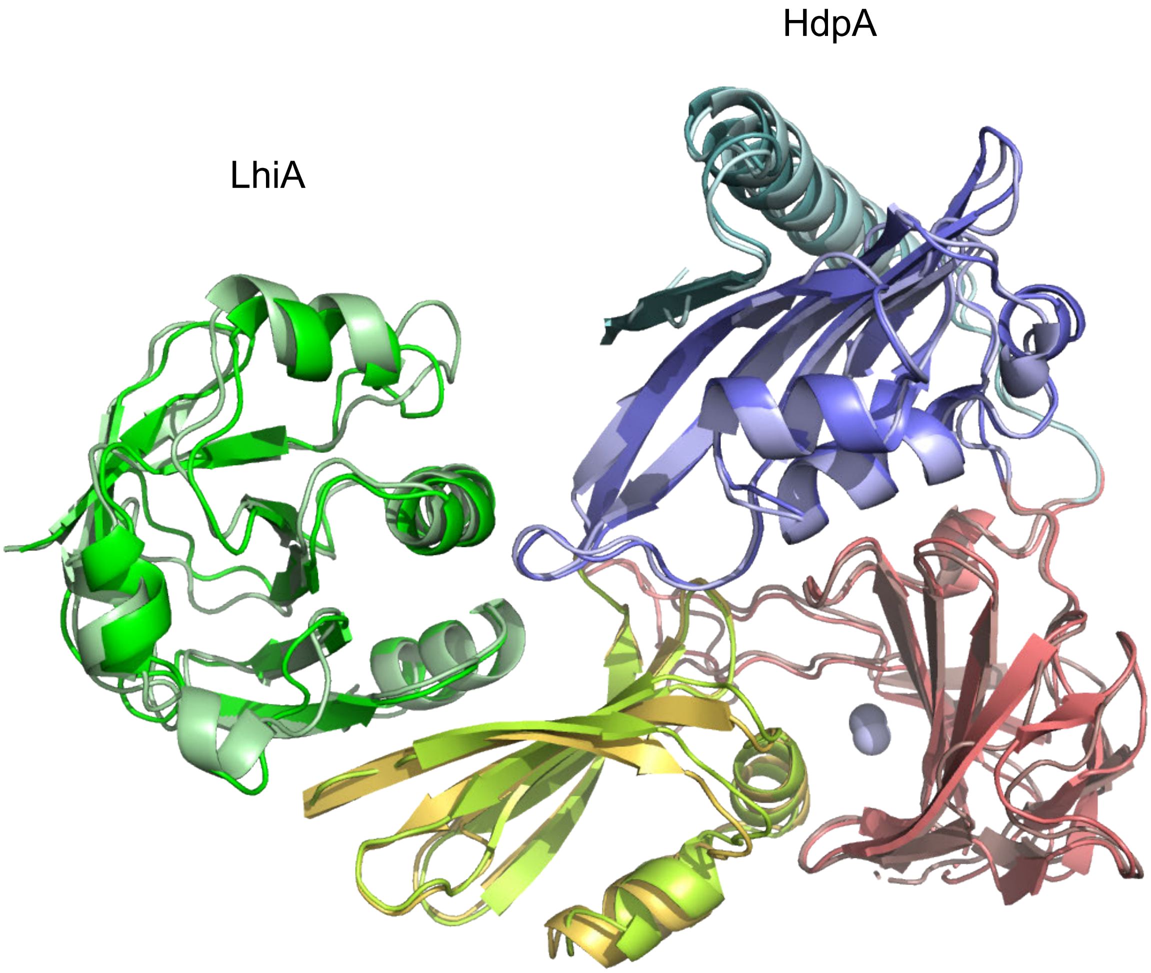


**Fig. S3.** The structure of the fusion LhiA$^{S}$-HdpA$^{S}$ and of the LhiA$^{S}$/HdpA$^{S}$ complex are similar. Structure of residues 45-185 of LhiA and 40-406 of HdpA within the fusion protein (same color code as Fig 2) were superimposed on the same residues of the complex (different shade of the same colors).

A

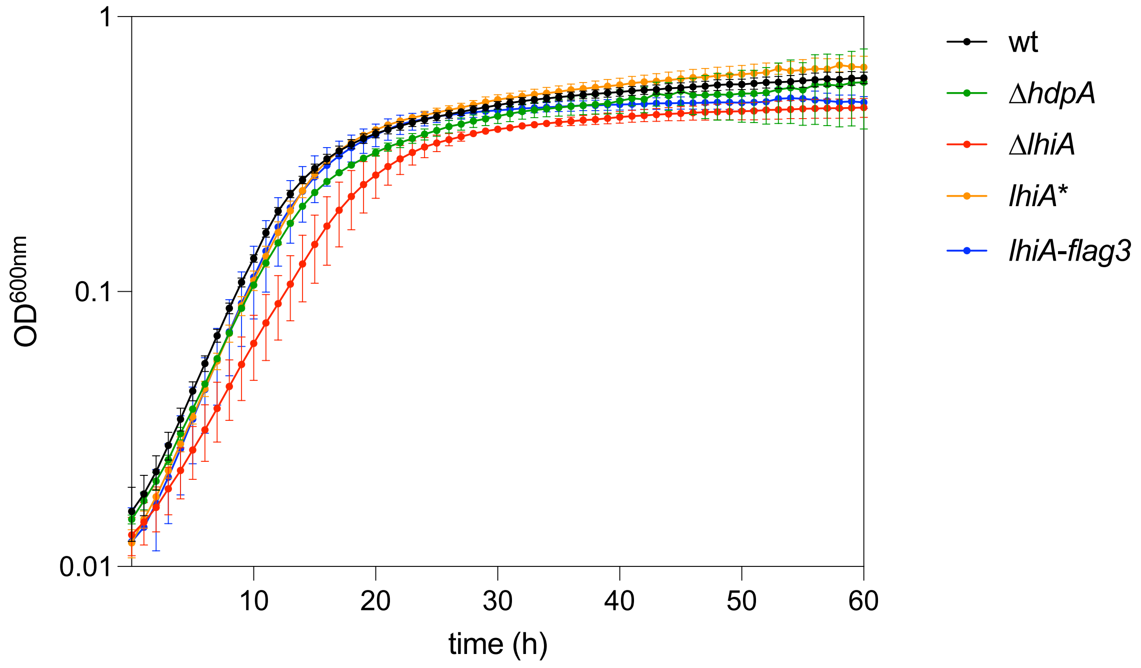


B

| relevant genotype | generation time (h) ± SD |
|---|---|
| wt | 3.09 ± 0.15 |
| *ΔhdpA* | 3.35 ± 0.15 |
| *ΔlhiA* | 4.00 ± 0.26 |
| *lhiA** | 3.06 ± 0.08 |
| *lhiA-flag3* | 2.89 ± 0.11 |

**Fig. S4.** Growth curves strains N6, N6 Δ*hdpA*::*aphA3*, OMD1, OMD2, OMD3. A. Cells from the indicated strains were grown at least in triplicate in 24-well plates (volume 1 mL) as indicated in the Material and Methods except that the $CO_2$ level was 10% in a Spark (TECAN) microplate reader. B. Average generation times are indicated.

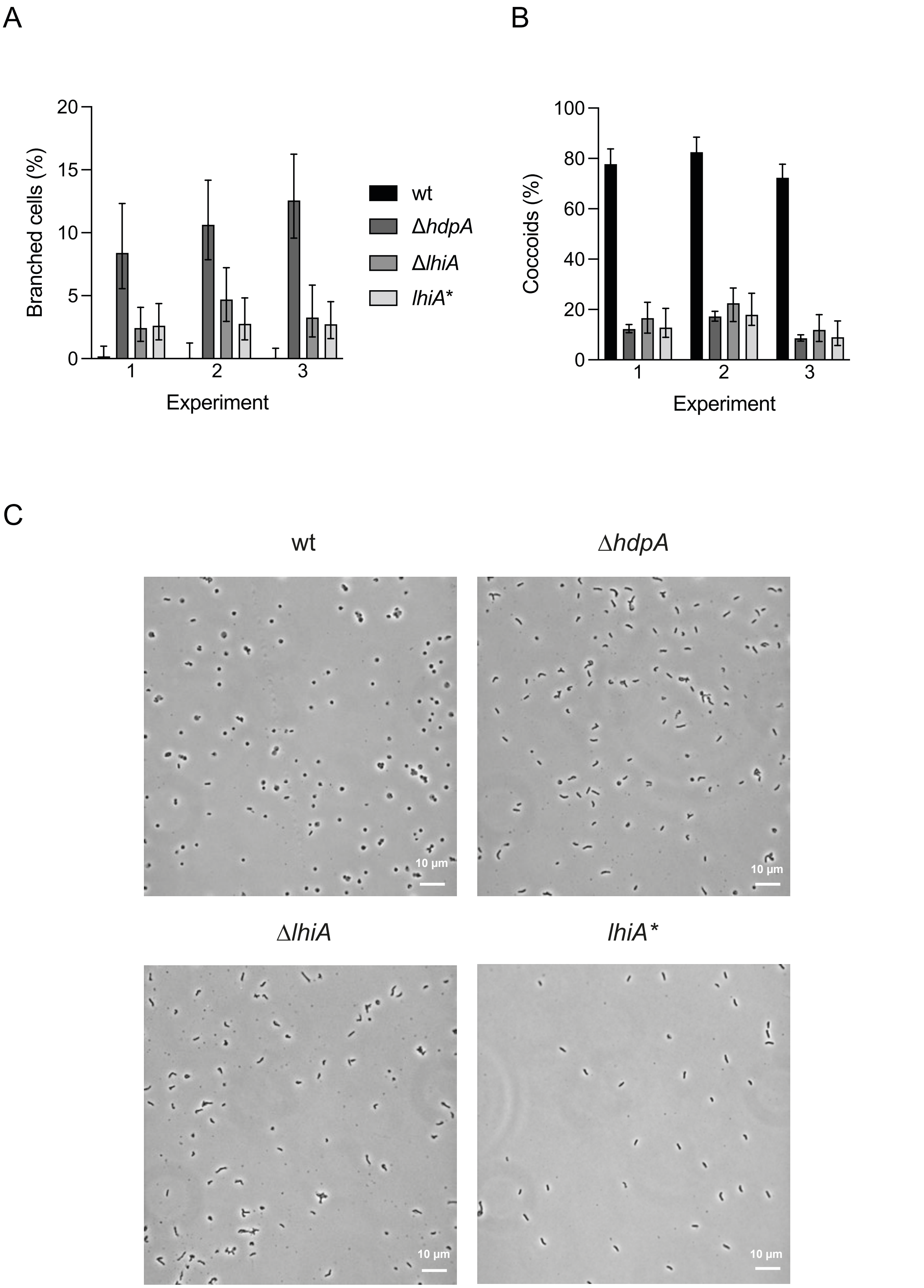


**Fig. S5.** A. Proportions of branched cells used in the plot of Fig. 4A. B. Proportions of coccoid cells used in the plot of Fig. 4B. Error bars represent the Wilson score 95% confidence intervals. Grayscale code as in A. C. Representative phase contrast images of cells from strains N6, N6 Δ*hdpA*::*aphA3*, OMD1, OMD2 after 120h growth.

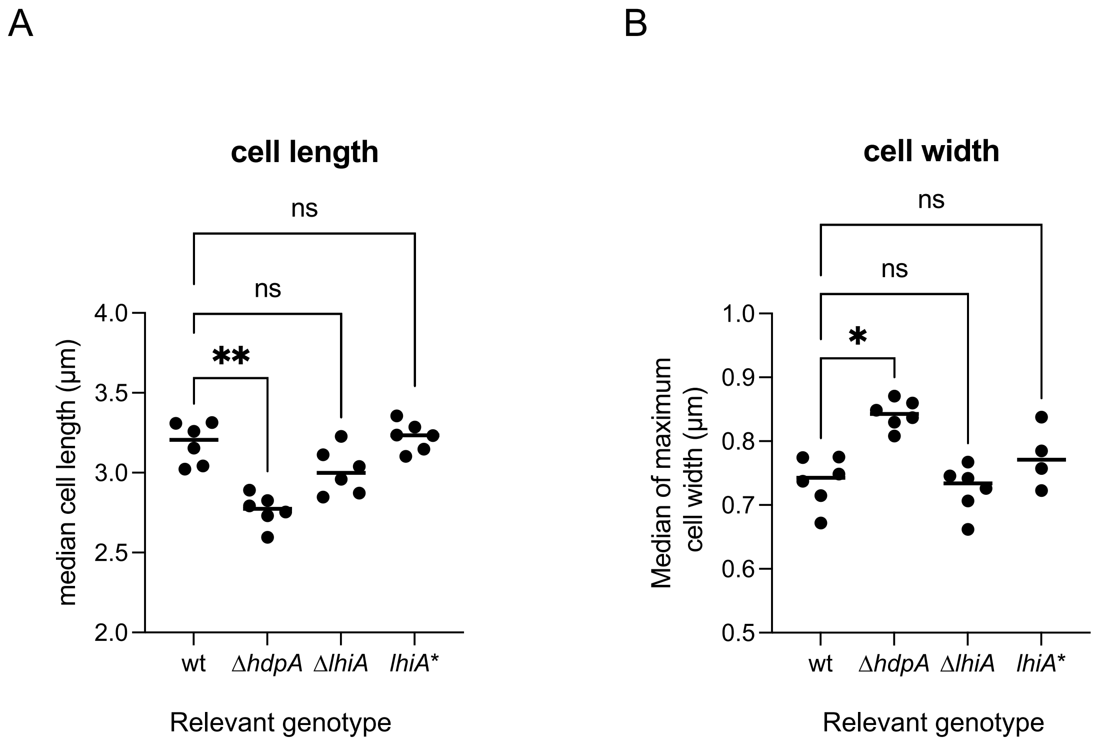


**Fig. S6.** Plots of the median cell width and cell length obtained from image analysis of phase contrast microscopy of four to six independent cultures of live cells from N6, N6 Δ*hdpA*::*aphA3*, OMD1, OMD2 at (OD 0.25-1.1). n>200 for each culture. Asterisks correspond to p-values of a Kruskal-Wallis test with correction for multiple comparisons.

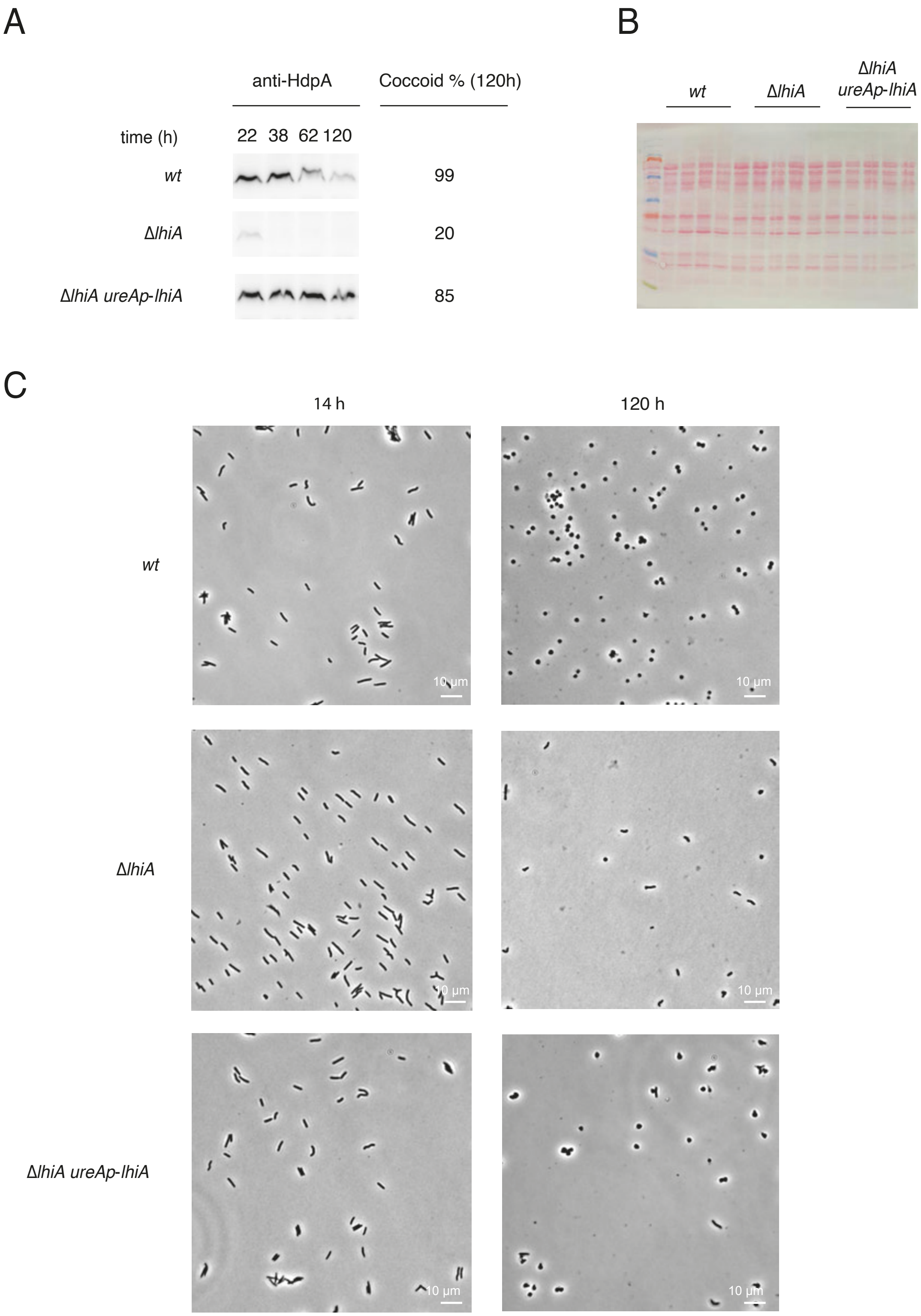


**Fig. S7.** Complementation of the Δ*lhiA* mutant. A. Complementation of the HdpA stabilization and coccoid transition phenotypes of the Δ*lhiA* mutation. Relevant genotypes are indicated. Western blots of SDS-PAGE resolved total extracts of strains with the indicated genotype were probed with anti-HdpA antibodies. Coccoid proportion among total cells at 120h were obtained from analysis of images of phase contrast microscopy on fixed cells (n>150). B. Ponceau staining of the membrane used for A. C. Representative microscopy phase contrast images of cells of the 4 strains.

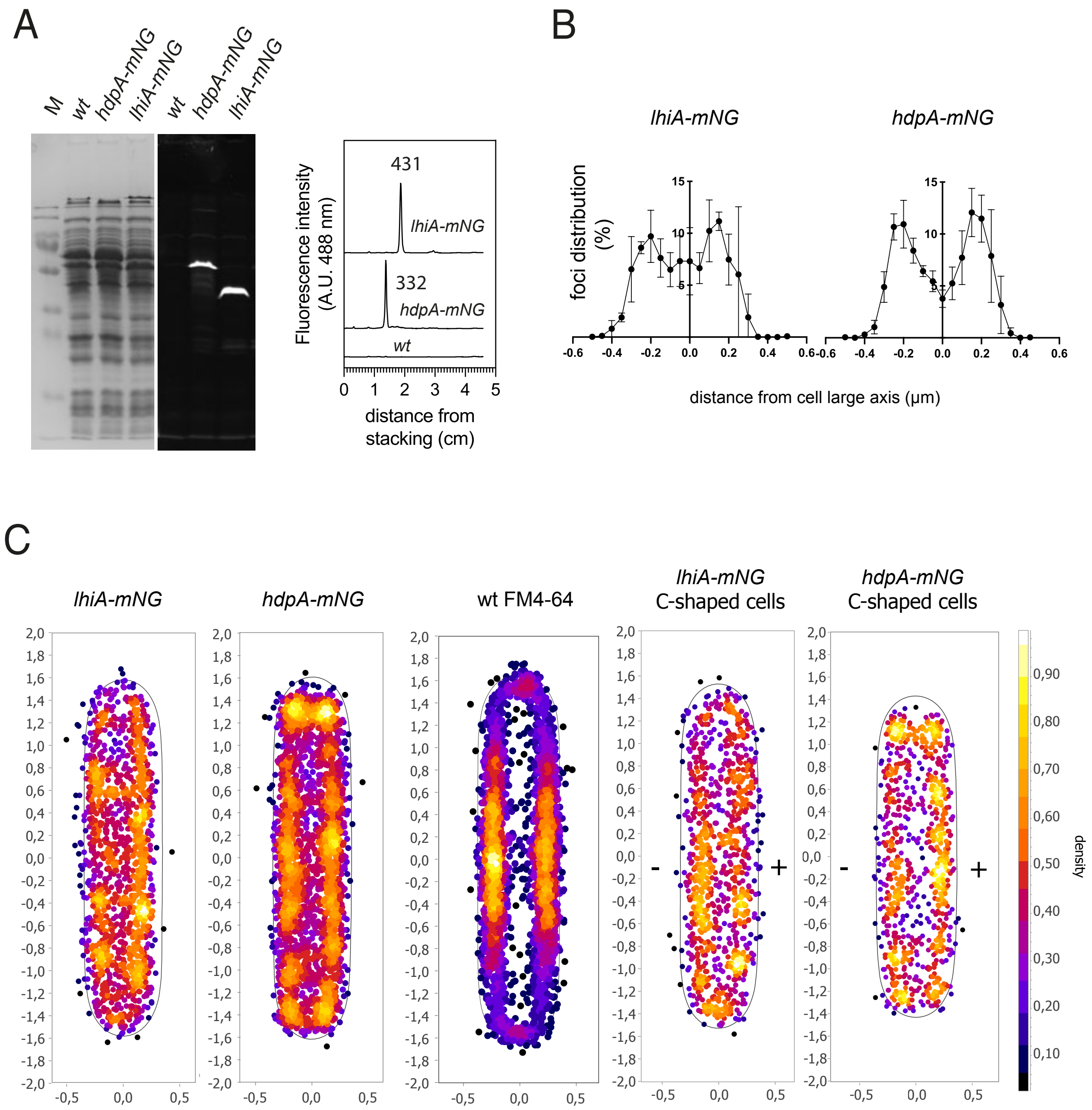


**Fig. S8.** Subcellular localization of LhiA and HdpA. A. SDS-PAGE analysis the HdpA-mNG fusion and LhiA-mNG proteins produced by strains OMD6 and OMD7. Left : early exponential phase (OD 0.16-0.20) total extracts of the two strains were run on a 13% gel which was directly imaged on a ChemiDoc™using Coomassie blue and Alexa 488 settings (A.U. arbitrary units). Right : quantification of the fluorescence. Plot profiles of gel lanes are shown with the relevant genotype of the strain used to generate the extract indicated. The numbers next to each peak correspond to the surface under the peak in arbitrary units. B Distribution of foci (fluorescence maxima, colored by cell density) along the transverse axis of the bacterial cells from strains OMD6, OMD7 and N6 in the presence of FM4-64 dye. Location of fluorescence foci was determined using MicrobeJ for more than 1000 cells for OMD6 and OMD7 and 778 for N6/FM4-64. C. representation of the density of the foci within an averaged straightened cell using the subcellular localisation tool of MicrobeJ. Curved cells were selected as described in the supplementary Material and Methods and the sides with apparent positive and negative curvatures are indicated by a + and a - sign, respectively.

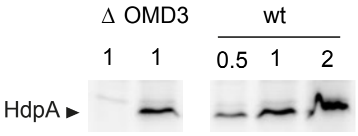


**Fig. S9.** Introduction of a 3xFlag encoding sequence at 3' end of the *lhiA* gene does not impair significantly LhiA ability to protect HdpA from proteolysis. Western blot of total extracts of strains N6Δ*hdpA::aphA3* (Δ), OMD3, and N6 (wt) grown in exponential phase (OD 0.42-0.58), probed with anti-HdpA antibodies. Numbers above the lanes correspond to the amount of material loaded relative to the OMD3 lane.

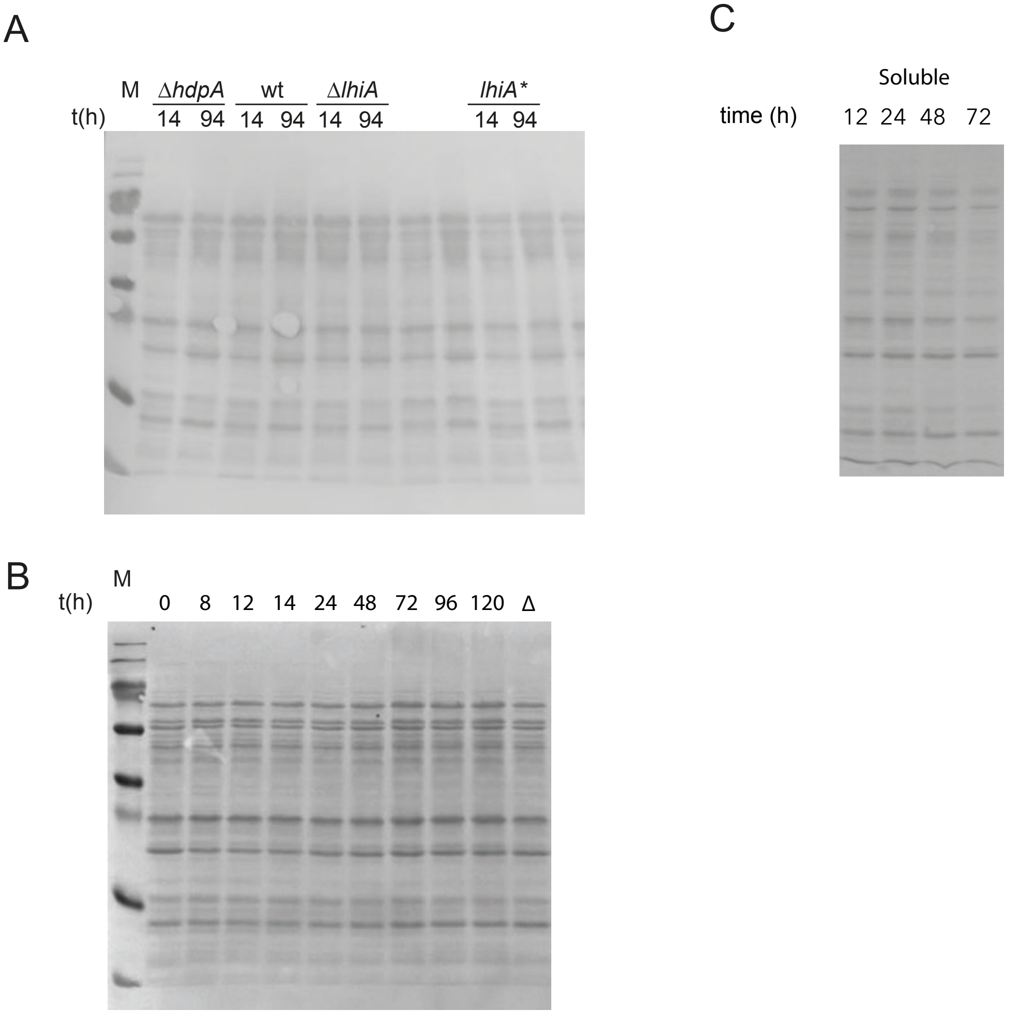


**Fig. S10.** Ponceau staining of the membranes used for the Western blots. A Membrane of Fig.5A. B. Total extract membrane of Fig. 6B. C. Soluble extract membrane of Fig. 6B.

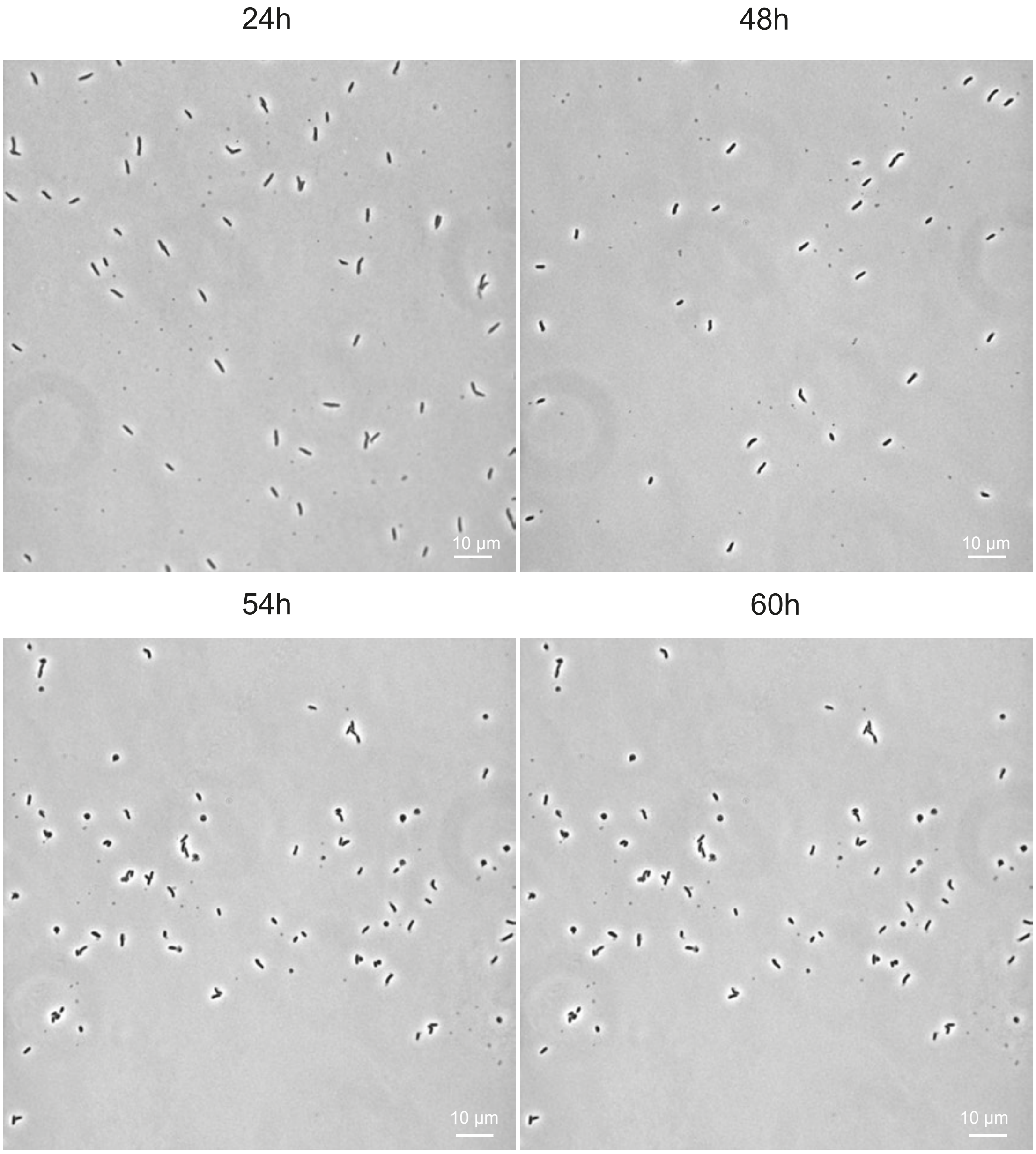
24h
48h
54h
60h
10 µm
10 µm
10 µm
10 µm

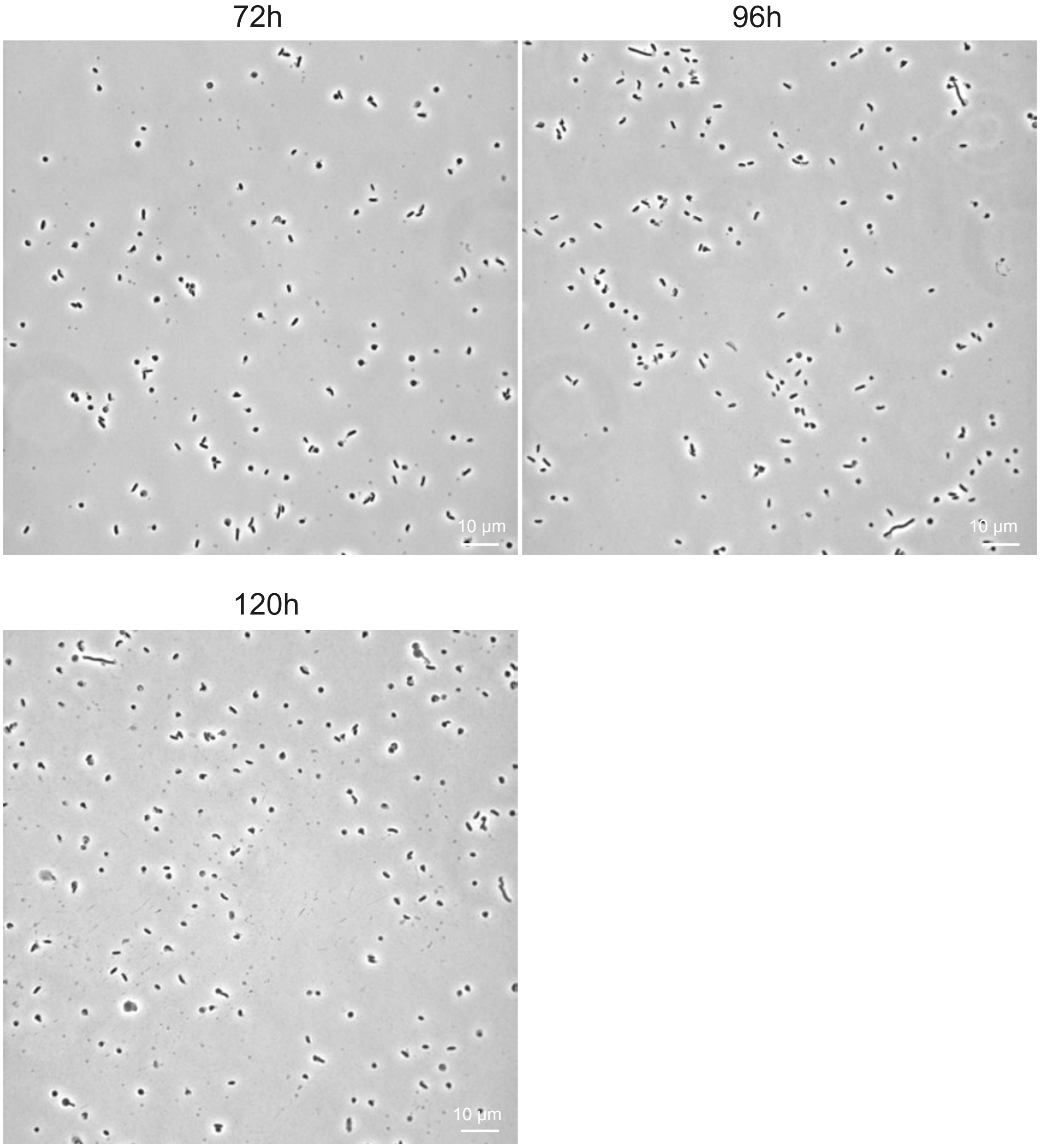


**Fig. S11.** Representative phase contrast microscopy images of OMD3 cells taken at different times (indicated above each image) for the coccoid quantification shown in Fig.6B.

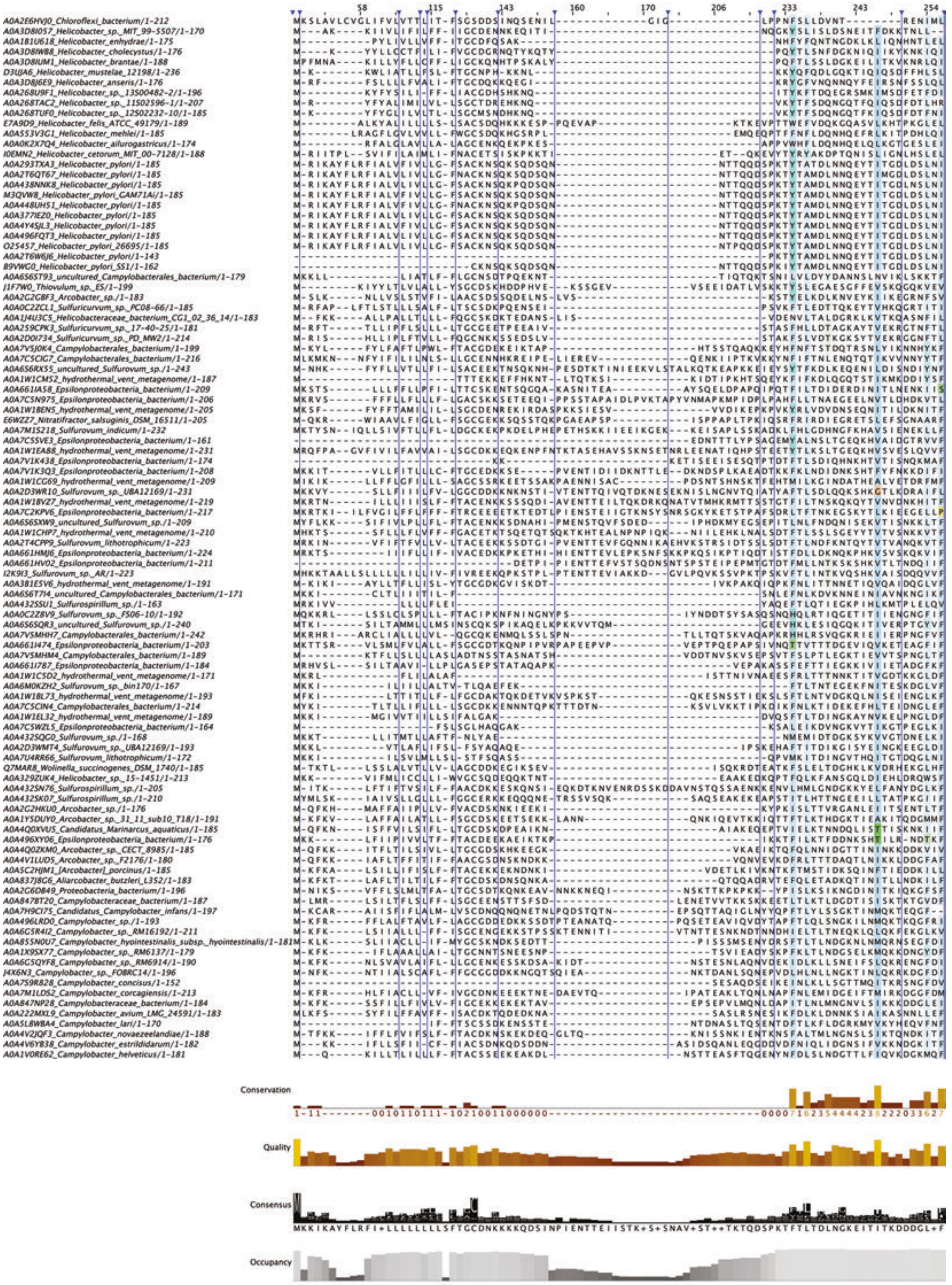
Conservation
Quality
Consensus
Occupancy

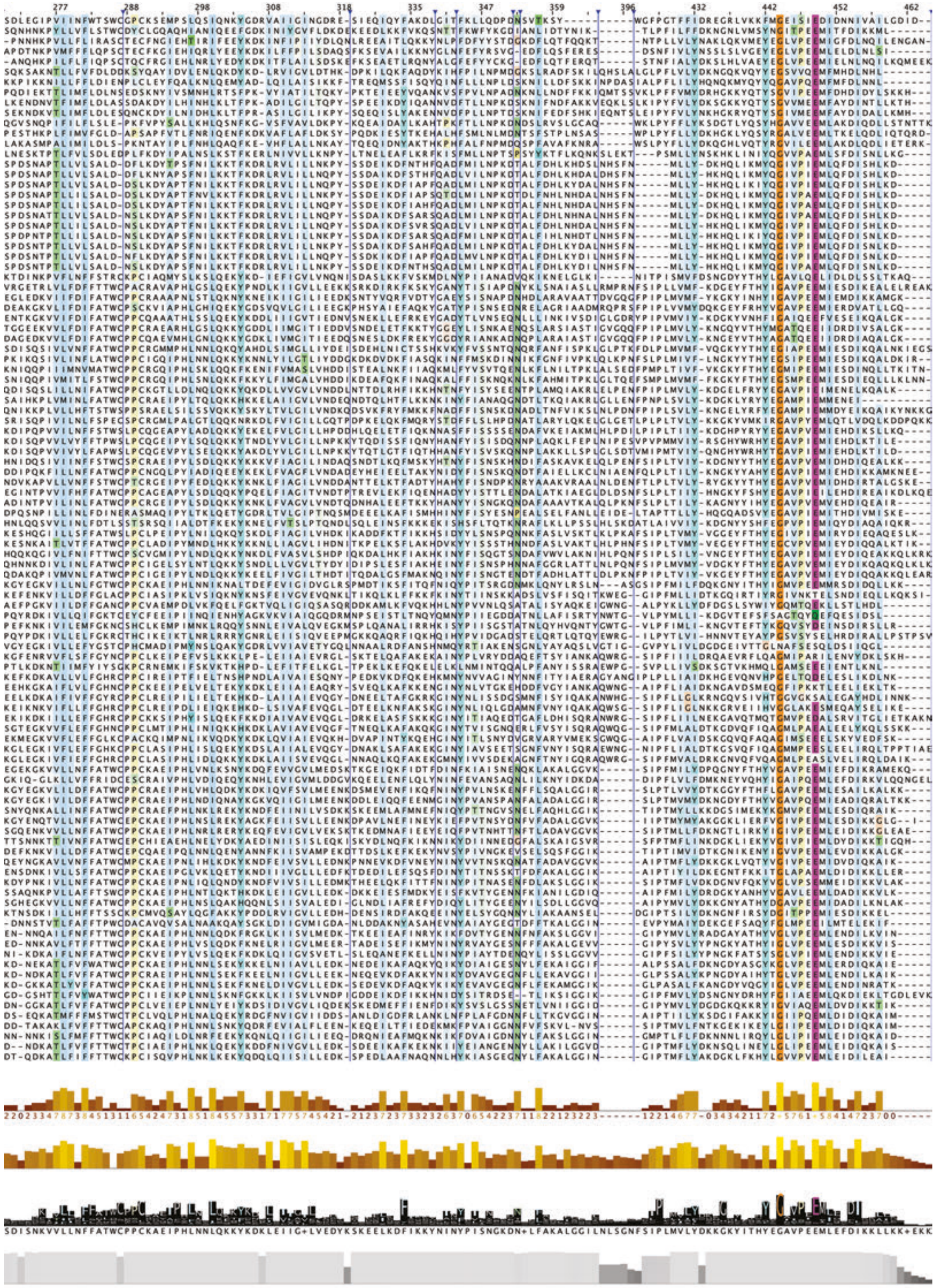

**Fig. S12.** MAFFT (5) alignment of the first 102 hits of a Foldseek search seeded with the Alphafold model of LhiA from H. pylori 26695, with color intensity determined by residue conservation. Jalview (6) was used for the visualization.

Campylobacter jejuni NCTC 11168

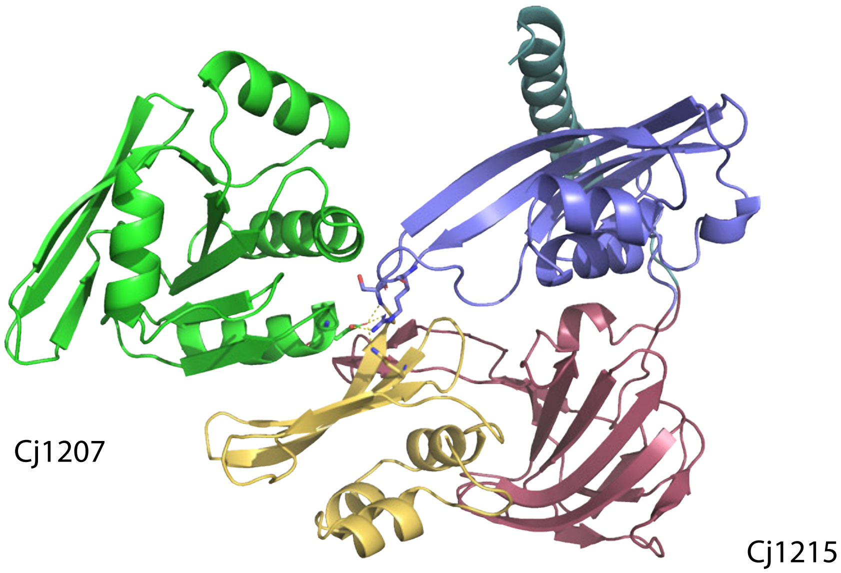


Nitratiruptor tergarcus DM16512

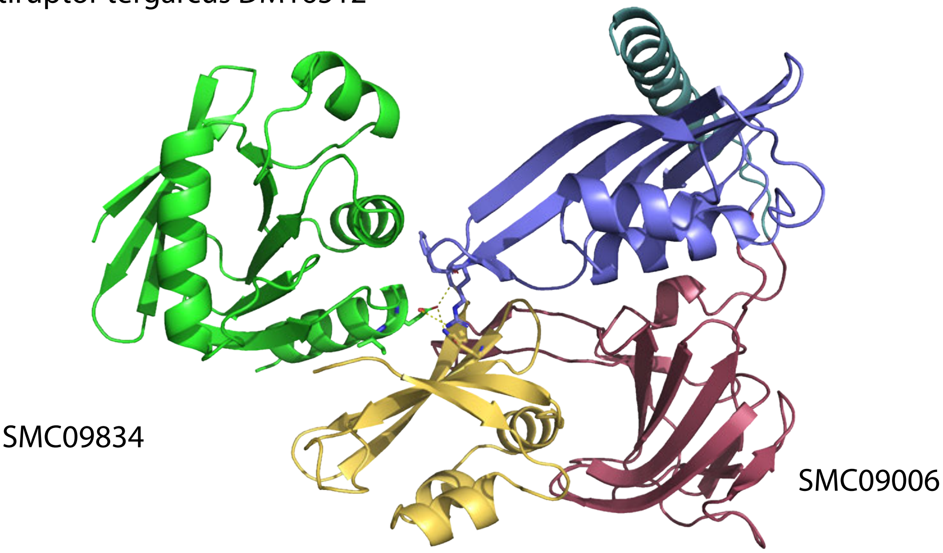


**Fig. S13.** Examples of Alphafold-2 generated models of complexes between the HdpA ortholog and the LhiA ortholog. The conserved interactions established by E174 with domain 1, domain 2 and the 1-2 linker are highlighted.

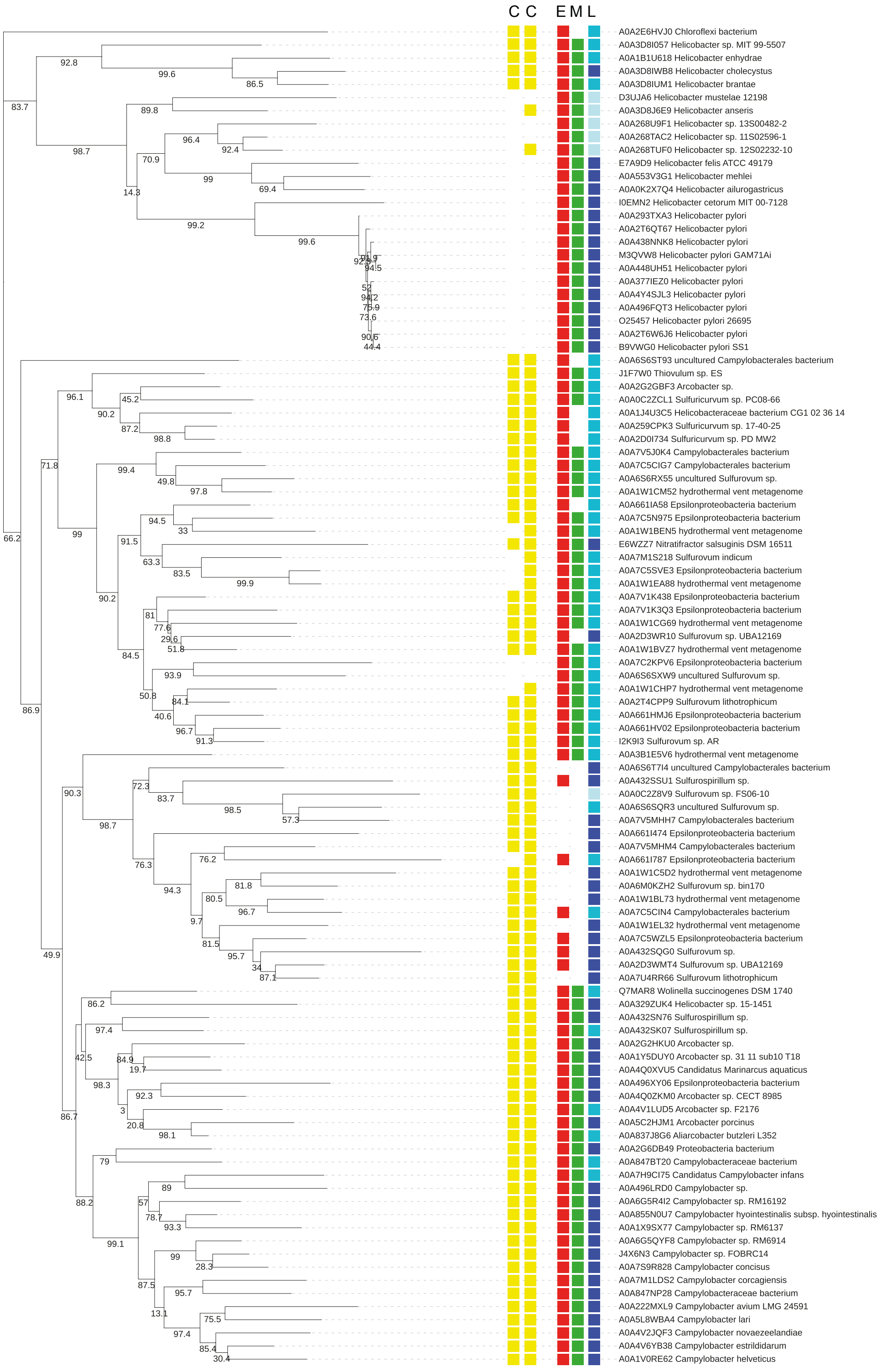


**Fig. S14.** Conservation of the thioredoxin cysteins and of the EML motif of the interface in the LhiA structural homologs. A Phylogenetic tree was generated from the alignment using iQTree (7) using the standard bootstrapping method and substitution model LG+F+G4 (5). The obtained tree was visualized using iTol (8), with bootstrap support indicated under each node. CC indicates presence (yellow boxes) or absence of the conserved cysteines of the thioredoxin fold. EML correspond to the conserved residues at positions 174 175 and 176 of LhiA. Position 174: presence of E is indicated by a red box, Position 175 : M, green. Position 176 L, dark blue, M, I, V : blue, F : light blue.

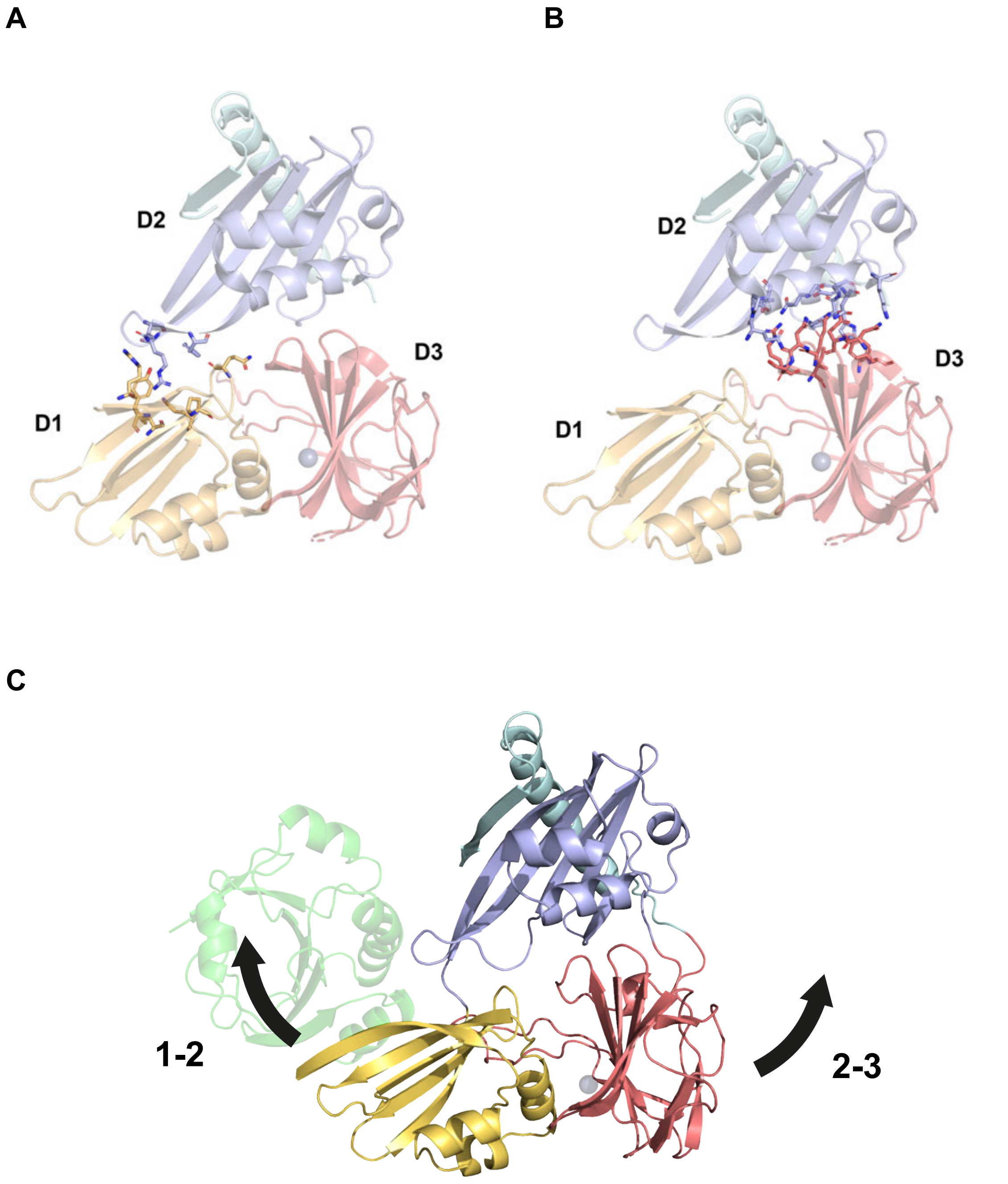


**Fig. S15.** Auto-inhibition and domain-domain interfaces of HdpA. A. Residues of the D1-D2 interface are highlighted (the D1-D2 linker has been removed for clarity) B. Residues of the D2-D3 interface are highlighted (D2-D3 linker removed). C. The two possible ways to relieve the HdpA auto-inhibition, shown on the LhiA-HdpA complex structure. Black arrows indicate possible domain motions that would relieve the occlusion of domain 3 by domain 1.

Table S1. **Data collection and refinement statistics for the LhiA-HdpA complex.**

| | **Danot_56** |
|---|---|
| **Wavelength** | 1.277 |
| **Resolution range** | 28.71 - 3.363 (3.62 - 3.36) |
| **Space group** | I 41 2 2 |
| **Unit cell** | 175.76 175.76 129.51 90 90 90 |
| **Total reflections** | 385016 (67560) |
| **Unique reflections** | 14581 (2747) |
| **Multiplicity** | 26.4 (24.6) |
| **Completeness (%)** | 99.09 (95.48) |
| **Mean I/sigma(I)** | 18.50 (2.64) |
| **Wilson B-factor** | 125.91 |
| **R-merge** | 0.1524 (1.322) |
| **R-meas** | 0.1554 (1.349) |
| **R-pim** | 0.02997 (0.2664) |
| **CC1/2** | 0.999 (0.792) |
| **CC*** | 1 (0.94) |
| **Reflections used in refinement** | 14553 (2747) |
| **Reflections used for R-free** | 727 (138) |
| **R-work** | 0.2332 |
| **R-free** | 0.2920 |
| **Number of non-hydrogen atoms** | 4044 |
| **macromolecules** | 4042 |
| **ligands** | 2 |
| **solvent** | 0 |
| **Protein residues** | 505 |
| **RMS(bonds)** | 0.010 |
| **RMS(angles)** | 1.49 |
| **Ramachandran favored (%)** | 89.18 |
| **Ramachandran allowed (%)** | 10.02 |
| **Ramachandran outliers (%)** | 0.80 |
| **Rotamer outliers (%)** | 9.32 |
| **Clashscore** | 7.03 |
| **Average B-factor** | 141.50 |
| **macromolecules** | 141.50 |
| **ligands** | 135.98 |

**Statistics for the highest-resolution shell are shown in parentheses.**

Table S2. **Data collection and refinement statistics for the LhiA-HdpA fusion protein.**

| | **Danot_65** |
|---|---|
| **Wavelength** | 0.9793 |
| **Resolution range** | 23.28 - 2.089 (2.277 - 2.089) |
| **Space group** | P 21 21 21 |
| **Unit cell** | 62.315 83.37 105.05 90 90 90 |
| **Total reflections** | 247901 (9810) |
| **Unique reflections** | 24207 (1210) |
| **Multiplicity** | 10.2 (8.1) |
| **Completeness (%)** | 91.8 (46.3) |
| **Mean I/sigma(I)** | 13.6 (1.5) |
| **Wilson B-factor** | 47.61 |
| **R-merge** | 0.095 (1.262) |
| **R-meas** | 0.100 (1.344) |
| **R-pim** | 0.031 (0.454) |
| **CC1/2** | 0.999 (0.732) |
| **Reflections used in refinement** | 24169 (264) |
| **Reflections used for R-free** | 1235 (16) |
| **R-work** | 0.2265 |
| **R-free** | 0.2597 |
| **Number of non-hydrogen atoms** | 4303 |
| **macromolecules** | 4096 |
| **ligands** | 1 |
| **solvent** | 206 |
| **Protein residues** | 505 |
| **RMS(bonds)** | 0.011 |
| **RMS(angles)** | 1.41 |
| **Ramachandran favored (%)** | 97.19 |
| **Ramachandran allowed (%)** | 2.61 |
| **Ramachandran outliers (%)** | 0.20 |
| **Rotamer outliers (%)** | 3.55 |
| **Clashscore** | 5.58 |
| **Average B-factor** | 53.17 |
| **macromolecules** | 53.14 |
| **ligands** | 51.70 |
| **solvent** | 53.72 |

**Statistics for the highest-resolution shell are shown in parentheses.**

**Table S3.** Summary of strain conctruction.

**Megaprimer constructs**

| | | L PCR | | | | R PCR | | |
|---|---|---|---|---|---|---|---|---|
| Strain | Genotype | L1 primer | L genomic DNA | L2 primer | Cassette | R1 | R genomic DNA | R2 |
| OMD1 | N6 Δ*lhiA*::*aphA3* | 762ΔG | N6 | 762KanG2 | *aphA*-3* | 762KanD3 | N6 | 762ΔD2 |
| OMD2 | N6 *lhiA* aphA3* | 762G_N6 | N6 | 762EMRR | *aphA*-3* | 762KanD3 | N6 | 762ΔD2 |
| OMD3 | N6 *lhiA-flag aphA3* | 762G_N6 | N6 | 762FlagKanG | *aphA*-3* | 762KanD3 | N6 | 762ΔD2 |

**Gibson constructs**

| Strain | Genotype | PCR A | | | PCR B | | | PCR C | | | PCR D | | |
|---|---|---|---|---|---|---|---|---|---|---|---|---|---|
| | | primer A1 | DNA A | primer A2 | B1 | DNA B | B2 | C1 | DNA C | C2 | primer D1 | DNA D | primer D2 |
| OMD5 | N6 Δ*lhiA*::*aphA3* Ω (*ureA*p-*lhiA* aac(3)IV) | 1499G | N6 | 73-1499R | 73-762F | N6 | Gibs 762-GentaD | Gibs 762-Genta G | N6*hp0772*::*aac(3)*IV** | Gibs Genta-1500 D | Gibs Genta-1500G | N6 | 1500D |
| OMD7 | N6 *lhiA*-mNeongreen-aphA3 | 762G_N6 | N6 | Neon-762 | 762-Neon | OMD 6 | KanRinv | Kan Grev | OMD1 | N6Δ762 | | | |

* cassette from pUC18K (1)

** Genomic DNA of a construct containing the sequence of the *aac(3)IV* from pUC1813apra (9).

**Table S4.** Summary of plasmid construction.

| | receiver | inserted DNA fragment | primers | cloning |
|---|---|---|---|---|
| pOD-1 | pET24b(+) NdeI-HindIII | HdpA$^S$ codons fused to LEHHHHHH encoding sequence, NdeI-HindIII | 762G, 762DH | ligation |
| pOD-2 | pET24b(+) NdeI-BamHI | LhiA$^S$ codons fused to HHHHHH encoding sequence, NdeI-BamHI | 506FP1, 506HRP1 | ligation |
| pOD-3 | pOD-1 NdeI | LhiA 25-185 codons, resulting in LhiA$^S$-HdpA$^S$ fusion with 3' LEHHHHHH encoding sequence, NdeI | pET762, 762-506hyb | ligation |
| pGEX-hdpA | pGEX-4T-1 BamHI-SmaI | *hdpA*, resulting in GST-hdpA fusion. | hdpABamHIF, hdpASmaIR | ligation |

**Table S5.** Criteria for the assignment of cells to rod vs coccoid.

| | Rods* | Coccoids |
|---|---|---|
| Area | 0.5-8 | 0.6-3 |
| Length | 1.5-5 | 0.5-2.7 |
| Width | 0.5-1.5 | 0.5-2.7 |
| Circularity | 0-max | 0.7-1 |

* When length/width <2, bacteria identified as rods were checked visually and reclassified as coccoid when they exhibited a spherical shape with one or two protruding poles.

**Table S6.** Primers used in this work

| name | sequence (5'-3') |
|---|---|
| 762ΔG | GTTGAATTCTACATGCCTAGCCCAAAACC |
| 762ΔD2 | GAGGATCCGATCGCGCTTTCTATGGTGT |
| 762KanG2 | GTTAGTCACCCGGGTACCGTATAAACACATCCTTAAGAATAATAAAATGGTAAC |
| 762KanD3 | CTATATTTTACTGGATGAATTGTTTTAGTACCTAAGGACTGAATTTAATGTTTAATTTTTTTAAAAAAATTG |
| 762FlagKanG | GTTAGTCACCCGGGTACCTCATTTATCGTCATCATCTTTGTAGTCCTTGTCATCATCGTCCTTATAGTCCTTATCGTCGTCATCCTTGTAATCCTTTAAGTTAGAAATATCAAATTGGAG |
| 762G_N6 | ATGAGAATTAAGGCTTATTTTTTGCG |
| 762G | GAACATATGCATCATCATCACCATCACAAAAAATCTCAAAAATCTCAAGATTCTCAAAAC |
| 762DH | GGGATCCTTAGTGATGGTGATGATGATGAGCGCTATCCTTTAAATTGGAAATATCAAATTG |
| 506FP1 | GAACATATGGCCGATGGAATGGCTAAAAAG |
| 506HRP1 | GAAAAGCTTTAGTGATGGTGATGATGATGTTCAAGAAAACCCTCTAAAAGATAAAATG |
| pET762 | GTTTAACTTTAAGAAGGAGATATACATATGAAAAATTCTCAAAAATCTCAAGATTCTC |
| 762-506hyb | GCCATTCCATCGGCACCGCTAGCGCTATCCTTTAAATTGGAAATATC |
| 762Flag-F | GATATTTCTAACTTAAAGGACTACAAGGATGACGACGATAAGGACTATAAGGACGATGATGACAAGGACTACAAAGATGATGACGATAAATGAGGATCCCCGGGTACCGAGCTCGAATTC |
| 762Flag-R | GAATTCGAGCTCGGTACCCGGGGATCCTCATTTATCGTCATCATCTTTGTAGTCCTTGTCATCATCGTCCTTATAGTCCTTATCGTCGTCATCCTTGTAGTCCTTTAAGTTAGAAATATC |
| 762N6_G | GAACATATGAGAATTAAGGCTTATTTTTTGCG |
| 762N6_D | GTAGGATCCTCAGTCCTTTAAGTTAGAAATATCAAATTG |
| 762EMRR-F26695 | AAGGGATCGTGCCAATAT |
| 762EMRR-Rev26695 | TATTGGCACGATCCCTTGATAC |
| pBR1 | TGACCAAAATCCCTTAACGTG |
| pBR2 | ACATTTCCCCGAAAAGTGC |
| pBR506 | CACGTTAAGGGATTTTGGTCAGGCATTAAACAACTGCATGG |
| pBR507 | GCACTTTTCGGGGAAATGTCCGGTGGTTTTAGCGTATTG |
| 506R2 | AAAACCCTCTAAAAGATAAAATGAATTTTTTTC |
| 507F2 | TAATGTCTTATTTTAAGAATGC |
| 507F3 | TAATGTCTTATTTTAAGAATGCTTTCAACC |
| 506-Neon | GAAAAAAATTCATTTTATCTTTTAGAGGGTTTTGGCGAGGAGGATAACATG |
| 507-Ne-sfT | GCATTCTTAAAATAAGACATTACTTGTACAGCTCGTCCATG |
| NeonR1 | CTTGTACAGCTCGTCCATGC |
| Neon-Kan | GACGAGCTGTACAAGTAATGGCTAAAATGAGAATATCACCG |
| Kan-507 | AGAGCTAGAACAAGGCTCTAAATACACACTAGAATCATCTATTAAAGATTTTTGGTTGAAAGCATTCTTAAAATAAGACATTAAAACAATTCATCCAGTAAAATATAGTATTTTATTTTC |
| 1499G | GTTGAATTCCATTACACCACCCATTGCAC |
| Gibs1499-762D | CGCTTGAAAAGGGCTAAAGCTGATTGGTTTAATCTCTGTATGAAGTC |
| Gibs1499-762G | TACAGAGATTAAACCAATCAGCTTTAGCCCTTTTCAAGCG |
| Gibs762-GentaD | CACGACTTTTATTCCTCCTATCAGTCCTTTAAGTTAGAAATATCAAATTG |
| Gibs762-GentaG | TTTCTAACTTAAAGGACTGATAGGAGGAATAAAAGTCGTGCAATAC |
| Gibs Genta- 1500D | TATTACAGAGCGAGATGGAATCAGCCAATCGACTGGC |
| Gibs Genta- 1500G | CTCGCCAGTCGATTGGCTGATTCCATCTCGCTCTGTAATAGG |
| 1500D | GAGGATCCCCCAAAGCAAACACCTTAGC |
| 73-762F | CATTGCTACAATTACGTCCAACCTTGATTTCGTTATGTCTTCAAGGAAAAACACTTTAAGAATAGGAGAATGAGATGAGAATTAAGGCTTATTTTTTGCG |
| 73-1499R | GTTGGACGTAATTGTAGCAATGTTTTGATTTACTAAGATTAACAGAAGCGTTCATGATTGGTTTAATCTCTGTATGAAGTC |
| 762_N6_FlagN_F | CAAGATTACAAAGATGATGACGATCAACAAGATAGCCCTAAAACCTAC |
| 762_N6_FlagN_R | TTGTTGATCGTCATCATCTTTGTAATCTTGAGATTTTTGAGATTTTTTACAAGC |
| 762wt-Kan | TATTTCTAACTTAAAGGACTGAGGTACCCGGGTGACTAAC |
| Kan-762wt | TTATTCCTCCTAGTTAGTCACCCGGGTACCTCAGTCCTTTAAGTTAGAAATATCAAATTGG |
| Neon-762 | TGGGAGAGAGGCCATGTTATCCTCCTCGCCGTCCTTTAAGTTAGAAATATCAAATTGGAG |
| 762-Neon | AGCAGAAATGCTCCAATTTGATATTTCTAACTTAAAGGACGGCGAGGAGGATAACATGGCCT |
| KanRinv | ACTCTTCCGAGCAAAGGACG |
| KanGRev | CGGTACCCGGGTGACTAAC |
| hdpABamHIF | CGCGGATCCGCCGATGGAATGGCTAAAAAG |
| hdpASmaIR | CCCGGGTTAAAAACCCTCTAAAAGATAAAATG |

.
